# Influence of Fluorination on Single-Molecule Unfolding and Rupture Pathways of a Mechanostable Protein Adhesion Complex

**DOI:** 10.1101/2020.07.09.194894

**Authors:** Byeongseon Yang, Haipei Liu, Zhaowei Liu, Regina Doenen, Michael A. Nash

## Abstract

Fluorination of proteins by cotranslational incorporation of non-canonical amino acids is a valuable tool for enhancing biophysical stability. Despite many prior studies investigating the effects of fluorination on equilibrium stability, its influence on non-equilibrium mechanical stability remains unknown. Here, we used single-molecule force spectroscopy (SMFS) with the atomic force microscope (AFM) to investigate the influence of fluorination on unfolding and unbinding pathways of a mechanically ultrastable bacterial adhesion complex. We assembled modular polyproteins comprising the tandem dyad XModule-Dockerin (XMod-Doc) bound to a globular Cohesin (Coh) domain. By applying tension across the binding interface, and quantifying single-molecule unfolding and rupture events, we mapped the energy landscapes governing the unfolding and unbinding reactions. We then used sense codon suppression to substitute trifluoroleucine (TFL) in place of canonical leucine (LEU) globally in XMod-Doc, or selectively within the Doc subdomain of a mutant XMod-Doc. Although TFL substitution thermally destabilized XMod-Doc, it had little effect on XMod-Doc:Coh binding affinity at equilibrium. When we mechanically dissociated global TFL-substituted XMod-Doc from Coh, we observed the emergence of a new unbinding pathway with a lower energy barrier. Counterintuitively, when fluorination was restricted to Doc, we observed mechano-stabilization of the non-fluorinated neighboring XMod domain. These results suggest that intramolecular deformation networks can be modulated by fluorination, and further highlight significant differences between equilibrium thermostability, where all constructs were destabilized, and non-equilibrium mechanostability, where XMod was strengthened. Future work is poised to investigate the influence of non-natural amino acids on mechanically-accelerated protein unfolding and unbinding reaction pathways.

## INTRODUCTION

Non-canonical amino acids (NCAAs) provide new chemical functionality in proteins, and can be used to modulate diverse properties including molecular recognition, stability, and activity.^1–3^ In particular, the unique chemical characteristics of fluorine can alter amino acid side chain properties including hydrophobicity, acidity, and reactivity.^4–6^ Incorporation of fluorinated analogues of aliphatic amino acids including leucine (LEU), isoleucine (ILE), and valine (VAL) imparts increased hydrophobicity^7–9^ and is typically marginally disruptive to native protein structure^10^. Isosteric hydrogen-to-fluorine replacement can therefore alter the protein folding energy landscape, in some cases increasing thermodynamic stability but in other cases weakening it.^11,12^ Incorporation of fluorinated amino acids^13–15^ has been shown to influence denaturation temperatures,^16–18^ binding affinity to specific ligands,^19^ catalytic enzyme activity,^20–22^ proteolytic resistance,^23,24^ and folding/aggregation propensity.^25^ Despite a wealth of knowledge on the influence of fluorinated amino acids on protein stability, one current knowledge gap is understanding how fluorination influences protein mechanical properties. To the best of our knowledge there are no prior studies investigating the influence of fluorination on protein mechanics.

Mechanical forces are fundamental in biology, and protein mechanical stability is an important non-equilibrium parameter that can influence behavior *in vivo.*^*26,27*^ Mechanical stability describes how much tension a folded domain can withstand prior to unfolding, or how much force is required to dissociate a receptor-ligand complex. Single-molecule force spectroscopy (SMFS)^28–31^ with the atomic force microscope (AFM) can be used to stretch single protein molecules, quantify intermediate folding states^32,33^ elucidate [un]folding energy landscapes while accounting for differences in loading geometry^34–38^ or the presence of dual modes of ligand recognition^39,40^. The goal of this work was therefore to investigate the role of protein fluorination on non-equilibrium mechanostability, specifically investigating any discrepancies in trends between equilibrium thermodynamic stability and non-equilibrium mechanostability.

As a model protein for this investigation, we chose a mechanostable Dockerin:Cohesin receptor-ligand complex from *Ruminococcus flavefaciens* (*Rf*). One binding partner in this complex comprises a tandem dyad of X-module (XMod) and Dockerin (Doc) that forms a non-covalent interaction with the other binding partner, Cohesin (Coh). We denote the complex XMod-Doc:Coh (**Figure 1A**) where the colon indicates the non-covalent binding interaction. This well-studied protein pair binds with nM affinity, and exhibits an O-ring^41^ binding patch comprising a hydrophobic center surrounded by hydrophilic polar and charged amino acids.^42^ Prior AFM-SMFS studies on this complex and close homologs^43–47^ have quantified two unfolding/unbinding reaction pathways and demonstrated that XMod stabilizes the Doc:Coh binding interface through an allosteric mechanism governed through contacts between XMod and the distal end of Doc, opposite its interface with Coh.^48–50^ As a consequence, when XMod is folded, the complex is activated and ruptures at high forces, whereas when XMod unfolds, the Doc:Coh interface is significantly weakened. This multi-domain polyprotein with interdependency in the mechanical properties therefore provided an opportunity to study the influence of fluorination on both XMod unfolding and Doc:Coh unbinding in the same experiment.

**Figure 1.**
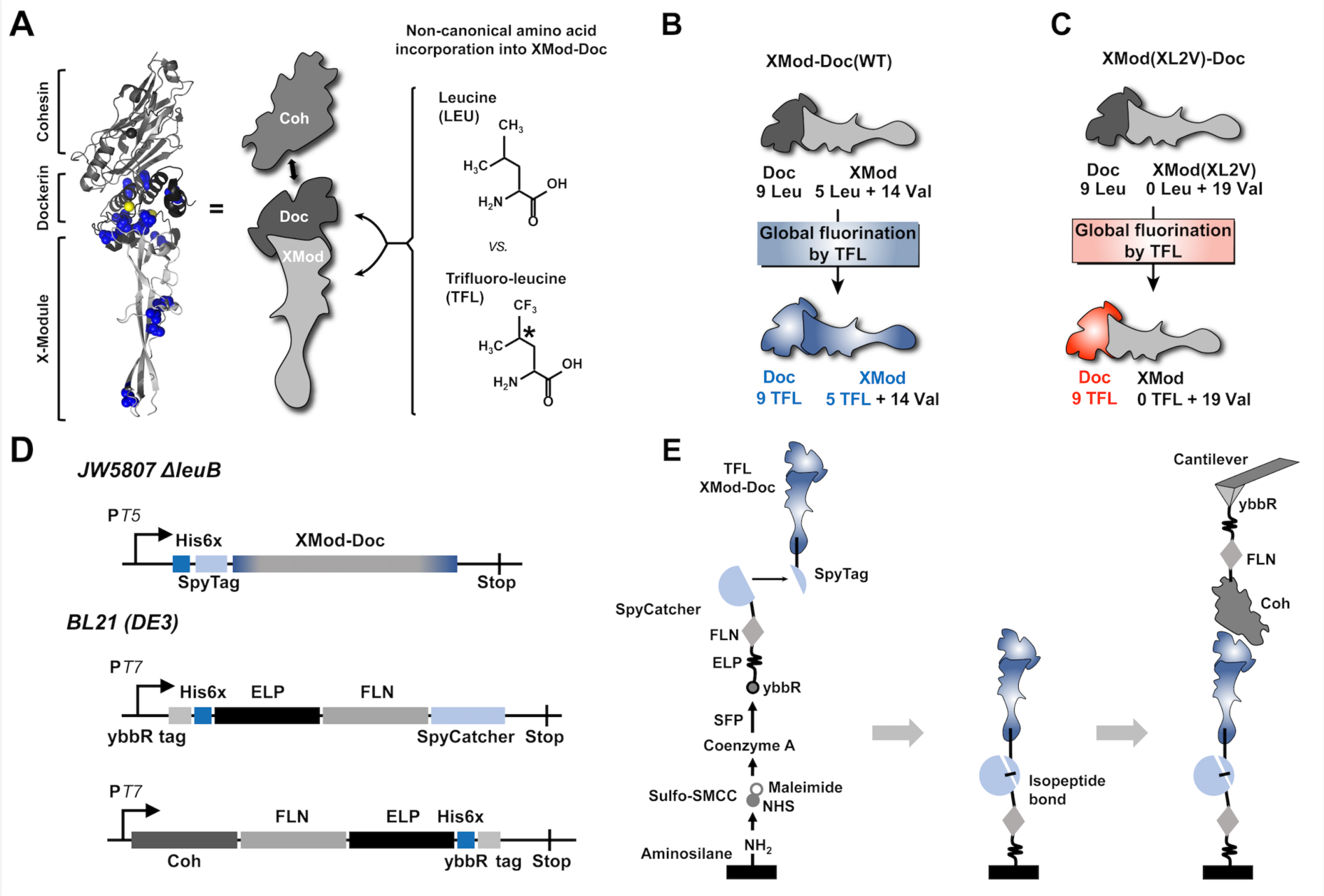
Design of XMod-Doc variants for domain-specific fluorination. (A) Crystal structure of XMod-Doc:Coh showing LEUs (blue), including 2 LEUs at the binding interface. Calcium ions are shown as yellow spheres. (PDB 4IU3). XMod-Doc is a single polypeptide chain containing sub-domains XMod and Doc that form a high-affinity non-covalent complex with Coh. The TFL structure shows the substituted –CF_3_ group in place of –CH_3_ in the side chain of LEU. The chiral center of TFL is indicated with an asterisk. (B) Global TFL incorporation into XMod-Doc (WT) replaced all LEU residues with TFL in both the XMod and Doc subdomains. (C) The mutant XMod(XL2V)-Doc was prepared by mutating all LEU codons in XMod to VAL. Global TFL incorporation resulted in fluorination only in Doc. (D) Gene cassettes for expression of TFL-incorporated His-SpyTag-XMod-Doc variants, linker protein ybbR-His-ELP-FLN-SpyCatcher, and Coh-FLN-ELP-His-ybbR. (E) Schematic illustration of surface chemistry and site-specific protein immobilization for AFM-SMFS.

We used TFL as the fluorinated amino acid, where a –CF_3_ group is substituted for one –CH_3_ methyl group in LEU. TFL is recognized by endogenous leucyl-tRNA synthetase (LeuRS), enabling quantitative replacement of LEU with TFL without requiring overexpression of endogenous or engineered LeuRS.^17,18,51^ We prepared wild type XMod-Doc (XMod-Doc (WT)) (**Figure 1B**) and two mutants where LEU residues in XMod or Doc were replaced with VAL, enabling localized incorporation of TFL into either the XMod or Doc sub-domains. We then analyzed WT and mutant protein complexes with canonical LEU or non-canonical TFL-incorporation using mass spectrometry, AFM-SMFS, thermal denaturation differential scanning fluorescence (DSF), and isothermal titration calorimetry (ITC) to quantify the effects of hydrogen-to-fluorine substitution in the hydrophobic side chain of LEU on mechanical and biophysical behavior.

## RESULTS AND DISCUSSION

### New mutant designs for localized fluorination and attachment chemistry

We designed XMod-Doc mutants by removing LEU codons from selected regions of the gene cassette encoding XMod-Doc, and replacing them with VAL codons. This allowed us to use global LEU sense codon suppression during expression runs in *E. coli* and achieve localized domain-specific incorporation of TFL into either XMod or Doc. The first new mutant is denoted XMod(XL2V)-Doc where all LEU codons from XMod were changed to VAL (**Figure 1C**). The second mutant was denoted XMod-Doc(DL2V) where all LEUs from Doc were mutated to VAL. For XMod-Doc(WT), residue-specific replacement of LEU by TFL resulted in TFL incorporation into both XMod and Doc subdomains. For XMod(XL2V)-Doc, TFL incorporation was localized to Doc, while for XMod-Doc(DL2V) TFL incorporation was localized to XMod. The WT and two mutant XMod-Doc proteins were produced using both canonical LEU and non-canonical TFL incorporation and thoroughly characterized to disentangle the influence of TFL on XMod unfolding and Doc:Coh unbinding.

To ensure we only analyzed valid single-molecule interactions, we performed AFM-SMFS with fingerprint domains^52–54^ attached to both the cantilever and surface molecules. We were concerned that TFL incorporation into fingerprint domains would change their unfolding signatures and severely reduce expression yields. To overcome this issue, we developed a novel scheme for site-specific immobilization of TFL-containing proteins for AFM-SMFS (**Figure 1D, E**). XMod-Doc variants (WT, XL2V, & DL2V) were designed and produced with an N-terminal SpyTag based on the plasmid pQE80L-SpyTag-ELP-SpyTag, a gift from Mark Howarth (Addgene plasmid # 112634 ; http://n2t.net/addgene:112634; RRID:Addgene_112634)^55^ (**Figure 1D**) which forms a spontaneous isopeptide bond with SpyCatcher^56,57^. We further produced a specialized surface-bound fingerprint protein (**Figure 1D**) containing SpyCatcher at the C-terminal end. This was fused at the DNA level with the fourth domain of *Dictyostelium discoideum* F-actin cross-linking filamin (FLN)^58^, which is an established unfolding fingerprint domain with a characteristic intermediate [un]folding state. In-frame with FLN-SpyCatcher on the N-terminal side, we included an elastin-like polypeptide (ELP) as an intrinsically disordered protein-based linker. The ELP allowed us to eliminate commonly used poly(ethylene) glycol linkers that are problematic in high-force protein unfolding studies due to mechano-isomerization of PEG that skews contour length analysis^59^. Finally, a ybbR/6x-histidine tag N-terminal to the ELP allowed for site-specific and covalent immobilization of the surface-bound fingerprint protein onto coenzyme A (CoA)-functionalized coverglass via ligation by 4’-phosphopantetheinyl transferase (SFP)^60^. Coh meanwhile was cloned as a single construct that had an FLN fingerprint domain, an ELP linker, and a C-terminal ybbR/6xHis tag. The gene cassettes are depicted in **Figure 1D**, and the surface chemical scheme is shown in **Figure 1E**.

### Incorporation of TFL

For TFL incorporation, we used the leucine auxotrophic strain *E. coli* ΔleuB (JW5807 from the Keio collection)^61^. Residue-specific incorporation was carried out using a standard media saturation method.^2,62^ After depleting a small amount of canonical leucine in minimal medium, the culture reached a stationary phase. We then supplemented the medium with TFL and induced expression of the target gene. TFL was charged onto leucyl-tRNAs by endogenous LeuRS and co-translationally introduced into XMod-Doc variants. Coh and SpyCatcher fusion proteins were expressed using a standard strain (BL21) in rich media (no TFL). All proteins were purified by metal ion affinity chromatography and size-exclusion. WT and variants expressed using LEU or TFL incorporation are denoted with the relevant amino acid appended after the protein abbreviation (e.g., XMod-Doc(WT)-LEU, XMod-Doc(WT)-TFL, XMod(XL2V)-Doc-LEU, XMod(XL2V)-Doc-TFL).

Sodium dodecyl sulfate polyacrylamide gel electrophoresis (SDS-PAGE) clearly showed successful expression and purification of TFL-incorporated XMod-Doc variants (**Figure 2A**). We measured TFL incorporation by high-resolution liquid chromatography electrospray ionization mass spectrometry (HRMS), and confirmed high yields of >92% for XMod-Doc(WT) and >90% for XMod(XL2V)-Doc (**Figure 2B,C**).

**Figure 2.**
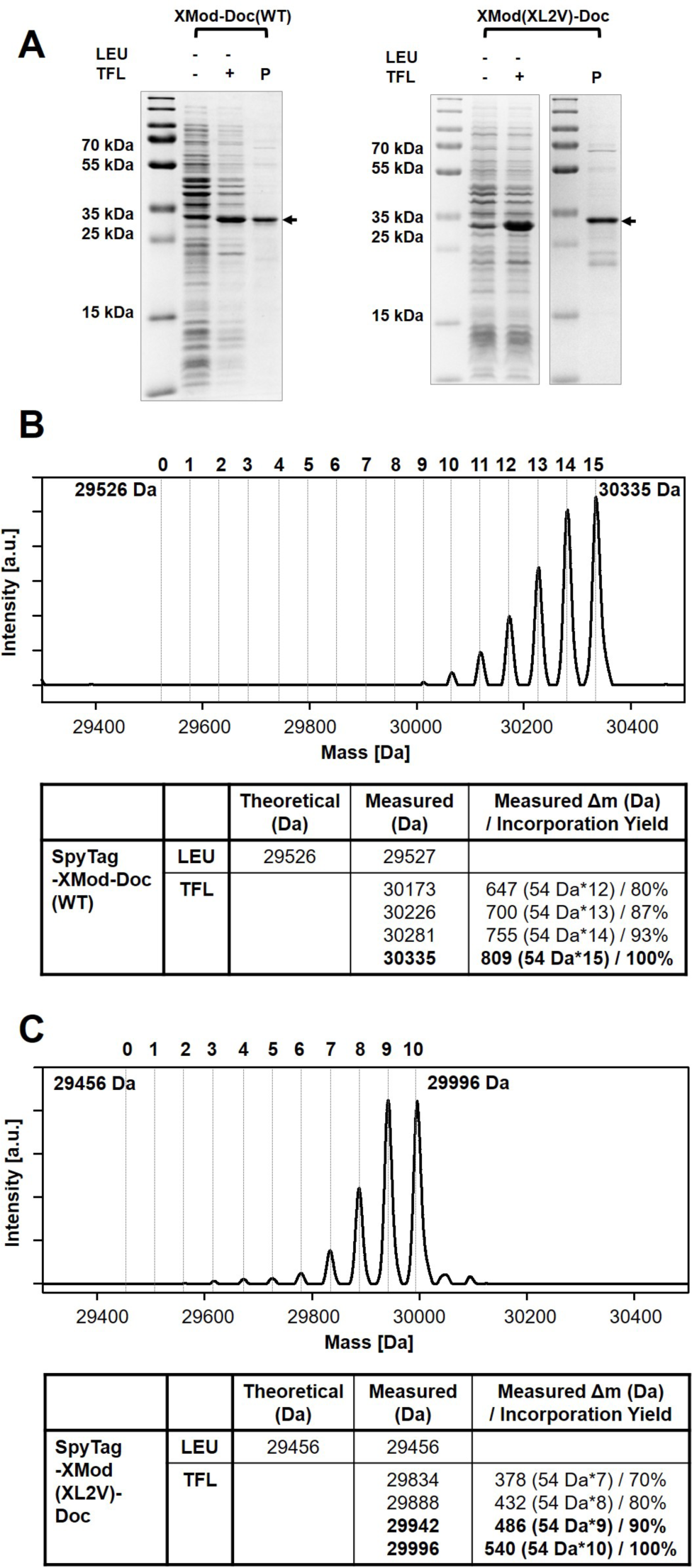
Residue-specific TFL incorporation into XMod-Doc. (A) SDS-PAGE analysis showing successful expression and purification of XMod-Doc(WT)-TFL and XMod(XL2V)-Doc-TFL. P: purified protein solution. (B) HRMS analysis of XMod-Doc(WT)-TFL. (C) HRMS analysis of XMod(XL2V)-Doc-TFL. Major mass peaks and corresponding incorporation yield of TFL are given in the tables.

Each successful TFL incorporation event is expected to increase protein mass by +54 Da due to the three hydrogen-to-fluorine substitutions (18 Da * 3). For XMod-Doc(WT)-TFL, the most intense peaks in the HRMS spectrum were 30,330 Da (with a mass difference of +54 Da*15 compared to XMod-Doc(WT)-LEU) and 30,281 Da (with a mass difference of +54 Da*14). These peaks corresponded to incorporation yields of 100% and 93%, respectively. For XMod(XL2V)-Doc-TFL, the most intense peaks in the HRMS spectrum were 29,942 Da (with a mass difference of +54 Da*9 compared to XMod-Doc(XL2V)-LEU) and 29,996 Da (with a mass difference of +54 Da*10), corresponding to incorporation yields of 90% and 100%, respectively. XMod-Doc(WT) and variants were also expressed with canonical LEU and characterized by SDS-PAGE and HRMS (**Figure S1, S2**). The expected molecular weights of Coh and the SpyCatcher fusion were validated by SDS-PAGE (**Figure S3**).

We tested the Coh-binding ability of XMod-Doc(WT), XMod(XL2V)-Doc, and XMod-Doc(DL2V) under canonical LEU incorporation using analytical size exclusion chromatography. XMod-Doc(WT)-LEU and XMod(XL2V)-Doc-LEU both bound Coh, however, XMod-Doc(DL2V)-LEU did not (**Figure S4**). This was not surprising given the two LEU residues at the hydrophobic center of Doc’s binding interface with Coh. Apparently the mutations from LEU to VAL in Doc were too deleterious to maintain binding ability. We terminated work on DL2V and no further results are reported on that variant. Additionally, we verified the functionality of SpyTag/SpyCatcher assembly using analytical chromatography (**Figure S5**).

### AFM-SMFS of XMod-Doc(WT)-LEU and -TFL

Next we performed receptor-ligand AFM-SMFS on XMod-Doc(WT)-LEU and XMod-Doc(WT)-TFL bound to Coh (**Figure 3A**). The surface was repeatedly probed with a Coh-modified cantilever tip, resulting in occasional formation of an XMod-Doc:Coh complex. The cantilever was retracted at constant speed, ELP linkers were stretched and tension built up in the system until sufficiently high forces were reached to unfold the two FLN fingerprint domains in series. Each FLN unfolding event contained an intermediate state, resulting in 4 low-force (∼60 pN) unfolding peaks. Data traces were filtered by searching for two 36 nm contour length increments with intermediate folding states associated with FLN unfolding (**Figure 3D,E**). FLN unfolding was followed either by rupture of the XMod-Doc:Coh directly (pathway 1 (P1), **Figure 3B**), or by unfolding of XMod and subsequent rupture of the Doc:Coh binding interface at greatly reduced forces (pathway 2 (P2), **Figure 3C**).

**Figure 3.**
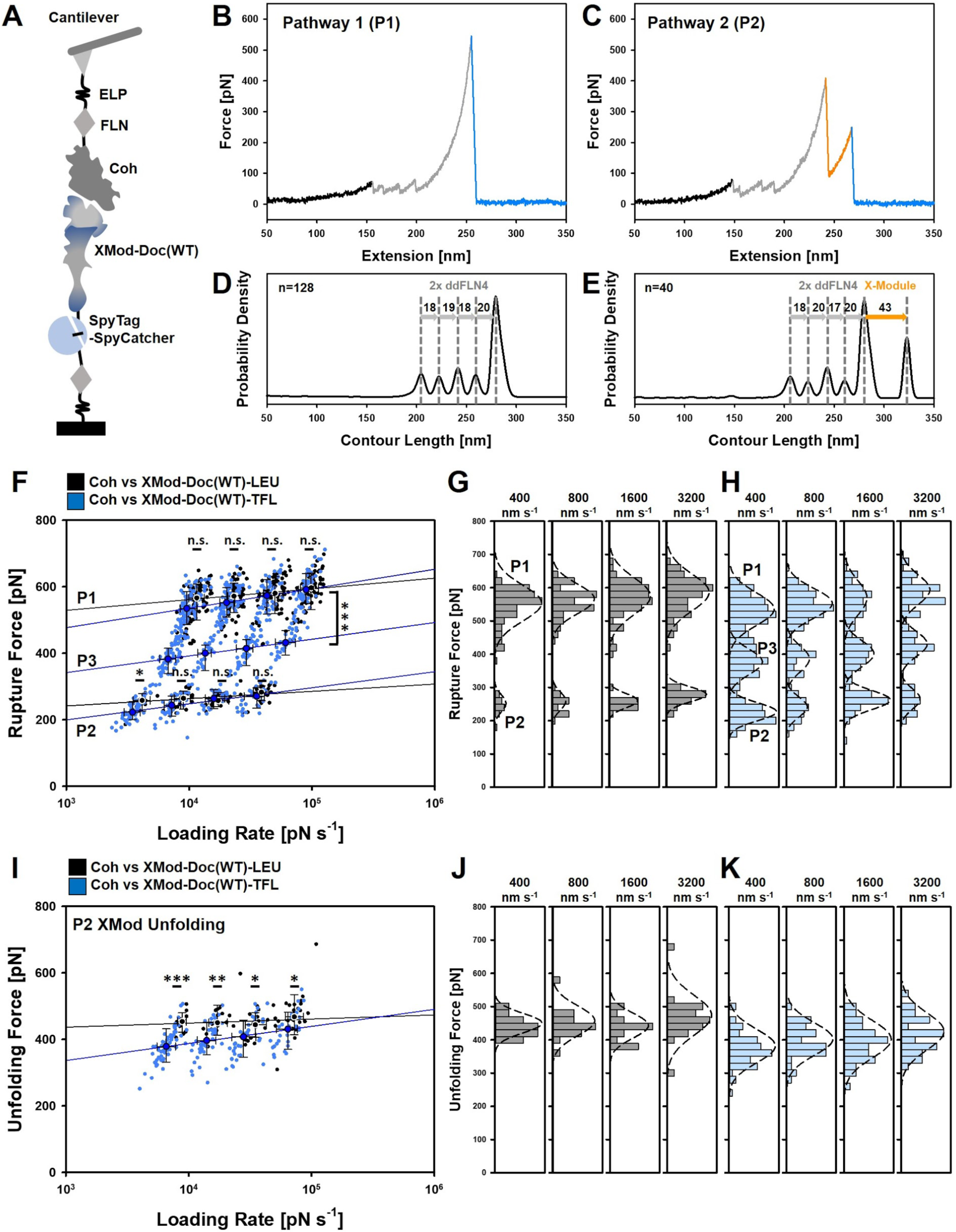
AFM-SMFS of XMod-Doc(WT)-LEU and -TFL. (A) Schematic illustration of experimental configuration. (B) Typical force-extension trace of XMod-Doc:Coh rupture with XMod remaining folded (pathway 1, P1). Unfolding of two FLN fingerprint domains (grey) was used to filter the curves for specific single-molecule interactions. Fingerprint unfolding was followed by complex rupture (blue). (C) Typical force-extension trace showing XMod unfolding followed by Doc:Coh rupture (pathway 2, P2). (D) Contour length histogram of P1 curves (n=128). (E) Contour length histogram of P2 curves (n=40). Increments between peaks were used for domain assignments to unfolding events. (F) Dynamic force spectra of XMod-Doc(WT)-LEU:Coh (black) and -TFL (blue) complex rupture forces for P1, P2 and P3 events. (G) Histograms of XMod-Doc(WT)-LEU:Coh P1 and P2 rupture events. (H) Histograms of XMod-Doc(WT)-TFL:Coh P1, P2, and P3 rupture events. (I) Dynamic force spectra of XMod unfolding events occurring along P2 pathways. (J) Histograms of XMod unfolding forces from XMod-Doc(WT)-LEU:Coh complexes. (K) Histograms of XMod unfolding forces from XMod-Doc(WT)-TFL:Coh complexes. Black and blue circles represent the median rupture force/loading rate at each pulling speed of 400, 800, 1600, and 3200 nm s^-1^. Error bars are ±1 s.d. Solid lines are least square fits to the Bell-Evans model. Statistical significance was determined with a t-test: n.s. *p* ≥ 0.01; * *p* < 0.01; \*\**p* < 0.001; and \*\*\**p* < 0.0001.

XMod-Doc:Coh complex rupture events from P1 and P2 pathways from filtered curves were analyzed and plotted as a function of the loading rate to generate a dynamic force spectrum (**Figure 3F**) and extract parameters that describe the free energy landscape of the unbinding reaction. Consistent with prior work^43,45^, XMod-Doc(WT)-LEU:Coh rupture events (**Figure 3F**, black dots) clearly showed two populations corresponding to a high-force population P1 (565-600 pN) and a low-force population P2 (255-285 pN). The same experiment was performed on XMod-Doc(WT)-TFL:Coh complexes. In addition to similar looking P1 and P2 rupture events, we observed a new rupture force population for the TFL-incorporated complex (**Figure 3F**, blue dots). This new rupture pathway, denoted P3, was situated at an intermediate force range of 380-435 pN between P1 and P2. The P3 pathway did not exhibit XMod unfolding prior to complex rupture, and was observed as a distinct population in the force histogram (**Figure 3H**).

We analyzed rupture events from P1, P2 and P3 rupture events obtained at several pulling speeds (400, 800, 1600, and 3200 nm s^-1^) and used the phenomenological Bell-Evans (BE)^63,64^ model as well as Dudko-Hummer-Szabo (DHS)^65,66^ theory to extract the energy landscape parameters, Δx, and *k*_*off*_, and additionally ΔG for the DHS model (**Table 1**). We found significantly lower Δx values for P1 and P2 rupture events for XMod-Doc(WT)-TFL:Coh rupture events as compared with XMod-Doc(WT)-LEU:Coh rupture events (**Table 1**). P3 events unique to TFL-incorporated complexes produced Δx values of similar magnitude to P1 of TFL-incorporated complexes, in a range of 1.6-2.0 Å (**Table 1**). We note that *k*_*off*_ fitting with the Bell-Evans model is not reliable enough to draw quantitative comparisons due to extreme model sensitivity^67^. Fitted Δx values, however, are generally robust. The observed trends in Δx (BE) and Δx/ΔG (DHS) for P1 and P2 rupture events indicate that global TFL replacement of LEU in XMod-Doc(WT) resulted in more rigid protein complexes with rupture events exhibiting a steeper loading rate dependency. The appearance of a distinct new P3 unbinding reaction pathway further indicates how TFL-incorporation modulates the energy landscape, in this case by enabling a new unbinding pathway with a lower barrier height.

**Table 1.** Energy landscape parameters of XMod-Doc:Coh complex rupture and X-module unfolding from AFM-SMFS from Bell-Evans (BE) model^63,64^ and Dudko-Hummer-Szabo (DHS) model^65,66^ (*v* = 0.5 and bin size of 20 pN).

| | | | $\Delta x$ (nm)<br>(BE) | $k_{off}$ (s<sup>-1</sup>)<br>(BE) | $\Delta G$ (k<sub>B</sub>T)<br>(DHS) | $\Delta x$ (nm)<br>(DHS) | $k_{off}$ (s<sup>-1</sup>)<br>(DHS) | $\Delta x/\Delta G$<br>(pm/k<sub>B</sub>T)<br>(DHS) |
| --- | --- | --- | --- | --- | --- | --- | --- | --- |
| **XMod-Doc(WT)** | **LEU** | **P1 (XMod folded)** | 0.30 | $2.9 \times 10^{-15}$ | 30.4 | 0.24 | $4.7 \times 10^{-8}$ | 7.8 |
| | | **P2 (XMod unfolded)** | 0.43 | $1.1 \times 10^{-9}$ | 25.4 | 0.31 | $6.2 \times 10^{-5}$ | 12.3 |
| | | **P2 XMod unfolding** | 0.78 | $3.1 \times 10^{-34}$ | 34.2 | 0.39 | $2.7 \times 10^{-10}$ | 11.3 |
| | **TFL** | **P1 (XMod folded)** | 0.16 | $2.8 \times 10^{-7}$ | 27.9 | 0.20 | $8.2 \times 10^{-7}$ | 7.2 |
| | | **P2 (XMod unfolded)** | 0.20 | $3.6 \times 10^{-3}$ | 21.9 | 0.26 | $1.0 \times 10^{-3}$ | 11.8 |
| | | **P3 (XMod folded)** | 0.19 | $7.2 \times 10^{-6}$ | 27.0 | 0.19 | $1.2 \times 10^{-4}$ | 7.1 |
| | | **P2 XMod unfolding** | 0.19 | $1.2 \times 10^{-5}$ | 23.3 | 0.20 | $1.0 \times 10^{-4}$ | 8.5 |
| **XMod (XL2V)-Doc** | **LEU** | **P1 (XMod folded)** | 0.19 | $2.0 \times 10^{-8}$ | 29.6 | 0.15 | $3.3 \times 10^{-5}$ | 5.1 |
| | | **P2 (XMod unfolded)** | 0.36 | $1.0 \times 10^{-6}$ | 26.2 | 0.34 | $6.8 \times 10^{-5}$ | 12.8 |
| | | **P2 XMod unfolding** | 0.20 | $2.9 \times 10^{-6}$ | 19.1 | 0.16 | $1.2 \times 10^{-3}$ | 8.4 |
| | **TFL** | **P1 (XMod folded)** | 0.28 | $1.1 \times 10^{-13}$ | 25.7 | 0.15 | $3.4 \times 10^{-5}$ | 6.0 |
| | | **P2 (XMod unfolded)** | 0.31 | $2.0 \times 10^{-5}$ | 20.0 | 0.36 | $1.6 \times 10^{-5}$ | 17.8 |
| | | **P2 XMod unfolding** | 0.32 | $1.9 \times 10^{-12}$ | 26.9 | 0.23 | $2.9 \times 10^{-6}$ | 8.6 |

In addition to unbinding/rupture events, we also analyzed XMod unfolding events obtained from TFL-incorporated and LEU-incorporated XMod-Doc(WT) (**Figure 3I,J,K**). XMod-Doc(WT)-TFL showed a significant decrease in the unfolding force of XMod (375-435 pN) compared to XMod-Doc(WT)-LEU (450-470 pN). Δx was significantly decreased from 0.78 nm (BE) and 0.39 nm (DHS) for XMod-Doc(WT)-LEU to 0.19 nm (BE) and 0.20 nm (DHS) for XMod-Doc(WT)-TFL. A similar trend was observed for the ratio of Δx/ΔG (DHS), which decreased from 11.3 pm/k_B_T for XMod-Doc(WT)-LEU to 8.5 pm/k_B_T for XMod-Doc(WT)-TFL (**Table 1**). Therefore, we found both for Doc:Coh unbinding as well as XMod unfolding, TFL-incorporation resulted in interaction/folding potentials that were shorter and more rigid for TFL-incorporated samples.

### AFM-SMFS of mutant XMod(XL2V)-Doc-LEU and -TFL

Next we analyzed XL2V mutants under both -LEU and -TFL incorporation (**Figure 4A**) using the same surface immobilization and measurement procedures. XL2V allowed TFL-incorporation to be localized to Doc while only canonical amino acids were present in XMod. XMod(XL2V)-Doc-LEU:Coh complexes (**Figure 4B**, gray) and XMod(XL2V)-Doc-TFL:Coh complexes (**Figure 4B**, red) both showed similar behavior for P1 rupture events, with XMod remaining folded and rupture occurring at 530-590 pN (-LEU) and 540-575 pN (-TFL). Δx values extracted from Bell-Evans fitting and Δx/ΔG values extracted from Dudko-Hummer-Szabo fitting of P1 rupture events showed that, in contrast to the trend observed in XMod-Doc(WT), TFL-incorporation in XMod(XL2V)-Doc resulted in binding potentials with a slightly flatter loading rate dependency (higher Δx and Δx/ΔG) as compared with XMod(XL2V)-Doc-LEU (Δx = 0.28 nm, Δx/ΔG = 6.0 pm/K_B_T, -TFL; Δx = 0.19 nm, Δx/ΔG = 5.1 pm/K_B_T, -LEU; **Table 1**). Therefore, while TFL-incorporation rigidified WT complexes, it may have rendered XL2V variants slightly more plastic in their deformation response, however the effect was small. P2 rupture events that occurred following XMod unfolding were in similar force ranges for XMod(XL2V)-Doc-LEU:Coh and XMod(XL2V)-Doc-TFL:Coh complexes (225-255 pN (-LEU); 215-250 pN (-TFL)), and also shared similar loading rate dependency with no statistically significant differences (**Figure 4B,C,D; Table 1**). The P3 pathway meanwhile that was observed in XMod-Doc(WT)-TFL:Coh complexes was absent in all XMod(XL2V)-Doc:Coh unbinding reactions. This finding implicates the significant role of XMod in stabilization of Doc:Coh interfaces in the alternative P3 pathway for XMod-Doc(WT)-TFL.

**Figure 4.**
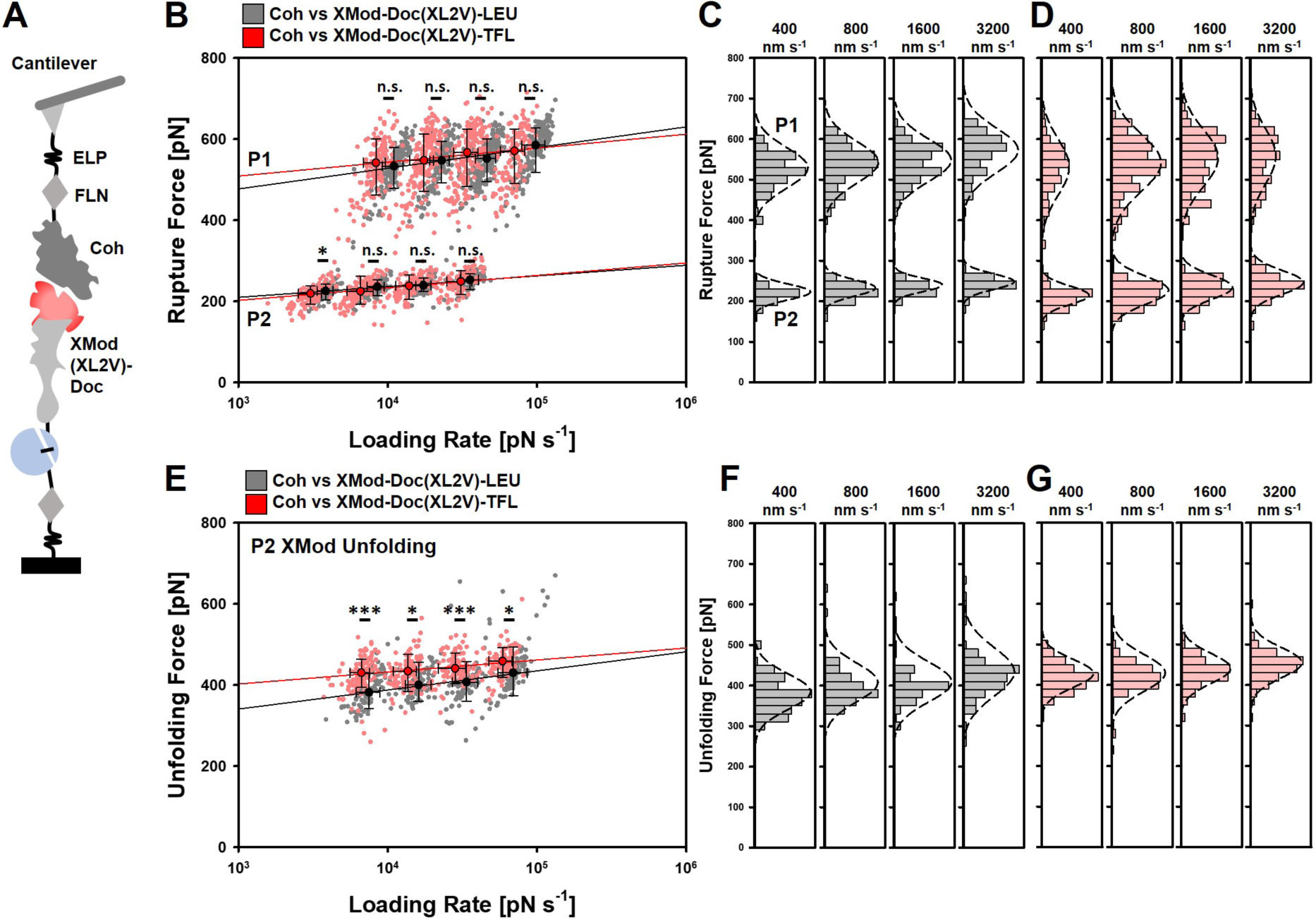
AFM-SMFS of XL2V variants under LEU- and TFL-incorporation. (A) Schematic illustration of experimental configuration, where TFL was incorporated into Doc but not XMod. (B) XMod(XL2V)-Doc-LEU:Coh (gray) and XMod(XL2V)-Doc-TFL:Coh (red) complex rupture events. (C) Histograms of XMod(XL2V)-Doc-LEU:Coh P1 and P2 rupture events at different pulling speeds. (D) Histograms of XMod(XL2V)-Doc-TFL:Coh P1 and P2 rupture event at different pulling speeds. (E) Dynamic force spectra of XMod unfolding events. (F) Histograms of XMod unfolding forces from XMod(XL2V)-Doc-LEU:Coh complexes at 4 pulling speeds. (G) Histograms of XMod unfolding forces from XMod(XL2V)-Doc-TFL:Coh complexes at 4 pulling speeds. Black and red circles represent the median rupture force/loading rate at each pulling speed of 400, 800, 1600, and 3200 nm s^-1^. All error bars are ±1 s.d. Solid lines are least square fits to the Bell-Evans model. Statistical significance was determined with a t-test: n.s. *p* ≥ 0.01; * *p* < 0.01; \*\**p* < 0.001; and \*\*\**p* < 0.0001.

We next analyzed XMod unfolding events occurring along the P2 rupture pathway for XMod(XL2V)-Doc-TFL and XMod(XL2V)-Doc-LEU at various pulling speeds, and extracted energy landscape parameters using BE and DHS fitting (**Figure 4E,F,G**). Interestingly, XMod(XL2V)-Doc:Coh complexes showed a significant increase XMod mechanical stability under TFL-incorporation as compared with LEU-incorporation (425-460 pN (-TFL); 380-435 pN (-LEU)). The Δx parameter showed a significant increase under -TFL incorporation (**Table 1**), again rendering the binding interface more compliant (Δx = 0.32 nm, Δx/ΔG = 8.6 pm/K_B_T, -TFL; Δx = 0.20 nm, Δx/ΔG = 8.4 pm/K_B_T, -LEU; **Table 1**) and reversing the trend observed upon TFL-incorporation into XMod-Doc(WT). TFL incorporation in Doc therefore significantly increased the force required to unfold the neighboring XMod domain.

We were concerned that small differences in cantilever spring constants could give rise to systematic errors on an absolute force scale. Therefore, to validate this result further, we performed AFM-SMFS experiments to probe XMod(XL2V)-Doc-TFL:Coh and XMod(XL2V)-Doc-LEU:Coh using the same cantilever (**Figure 5**). Both proteins were immobilized at different positions on the same glass slide, and a single Coh-modified cantilever was used to alternatively pick up and stretch molecules at each location at a pulling speed of 100 nm s^-1^. This experiment eliminated uncertainties based on cantilever calibration error or differences in extension values that arise because molecules are attached at different heights onto the AFM tip. The results confirmed that the XMod was slightly stabilized in XMod(XL2V)-Doc-TFL:Coh complexes and required higher forces to mechanically unfold.

**Figure 5.**
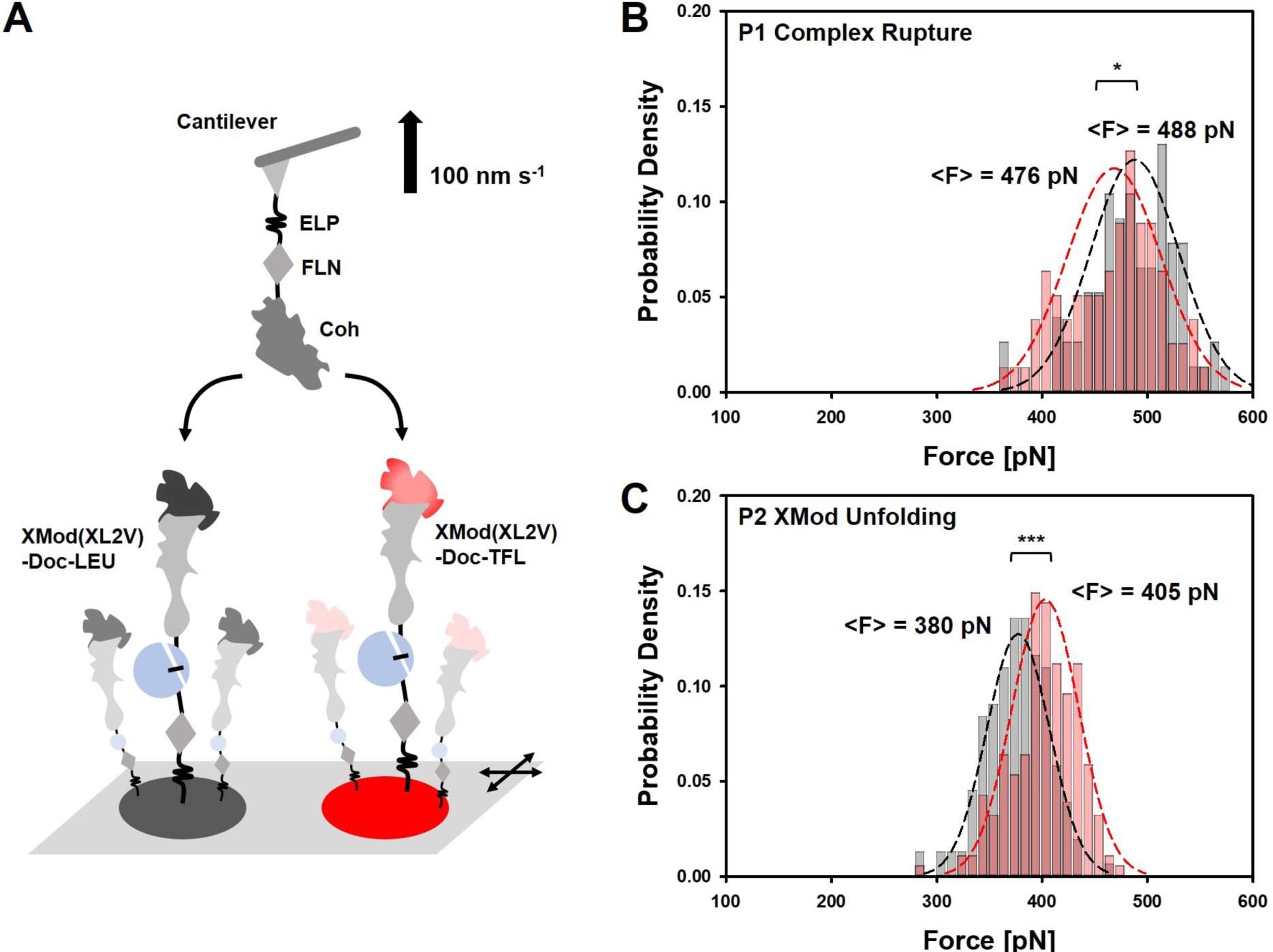
Comparative AFM-SMFS of XL2V variants under LEU- and TFL-incorporation. (A) Schematic of the experimental setup. XMod(XL2V)-Doc-LEU and -TFL were immobilized on different spots on the surface. Coh was immobilized at the cantilever tip. The same cantilever was used to alternate between two sample spots and probe each sample intermittently at 100 nm s^-1^. (B) Histograms of XMod(XL2V)-Doc-LEU and -TFL:Coh P1 rupture event. (C) Histograms of XMod(XL2V)-Doc-LEU and -TFL:Coh P2 XMod unfolding event. Statistical significance was determined with a t-test: n.s. *p* ≥ 0.01; \**p* < 0.01; \*\**p* < 0.001; and \*\*\**p* < 0.0001.

It could be argued that the higher XMod unfolding forces are observed by a ceiling or biasing effect^68^ of the receptor-ligand complex. A higher complex stability, one could argue, would result in higher unfolding force distributions for XMod unfolding events simply because the handle used to pull on XMod is stronger and the unfolding/rupture force distributions are overlapping. However, we verified that P1 and P2 Doc:Coh stability was nearly identical for the XL2V mutants under LEU- and TFL-incorporation (**Figure 4B, Figure 5B**). In fact, P1 complex rupture events for XMod(XL2V)-Doc-TFL:Coh were even slightly lower than those for -LEU, therefore, the mechanical strengthening we observe for XMod in -TFL complexes would in fact be *underestimated* by the biasing effect. The observed differences in XMod stability are therefore not explained by a statistical biasing effect of the complex on the XMod unfolding force distribution. We can speculate that an increase in hydrophobicity within Doc upon TFL-incorporation influenced the contact interface between Doc and XMod. The modified contacts could reasonably reroute the force propagation pathway^44–46^ through the molecule, introducing propagation paths with components perpendicular to the pulling axis. Such a scenario would result in an XMod domain that is more effective at distributing the force across its cross section, thereby requiring higher tension to unfold. By contrast, recall that global TFL incorporation into XMod-Doc(WT) significantly decreased the unfolding force of XMod (**Figure 3H**). In depth analysis by steered molecular dynamics simulation might offer the insight into the mechanism by which fluorination can alter these force propagation pathways.

### Bulk biophysical properties stability by ITC and NanoDSF

To investigate the effects of TFL incorporation on equilibrium binding affinity, we performed isothermal titration calorimetry (ITC) (**Figure 6A**). Analysis of the XMod-Doc(WT)-LEU:Coh interaction revealed K_D_ = 60 nM (K_D_ range (±σ) = 27-121 nM), while the XMod-Doc(WT)-TFL:Coh interaction showed K_D_ = 96 nM (K_D_ range (±σ) = 52-170 nM). Analysis of XMod(XL2V)-Doc-LEU:Coh resulted in K_D_ = 123 nM (K_D_ range (±σ) = 70-212 nM), while that of XMod(XL2V)-Doc-TFL:Coh showed K_D_ = 114 nM (K_D_ range (±σ) = 62-201 nM). Global LEU to TFL substitutions in the WT sequence may have slightly destabilized the interaction at equilibrium, however given the uncertainties associated with the method these observed differences are not considered significant. Meanwhile, LEU to VAL mutations in Doc (XMod-Doc(DL2V)-LEU) abolished binding ability completely (**Figure S4**).

**Figure 6.**
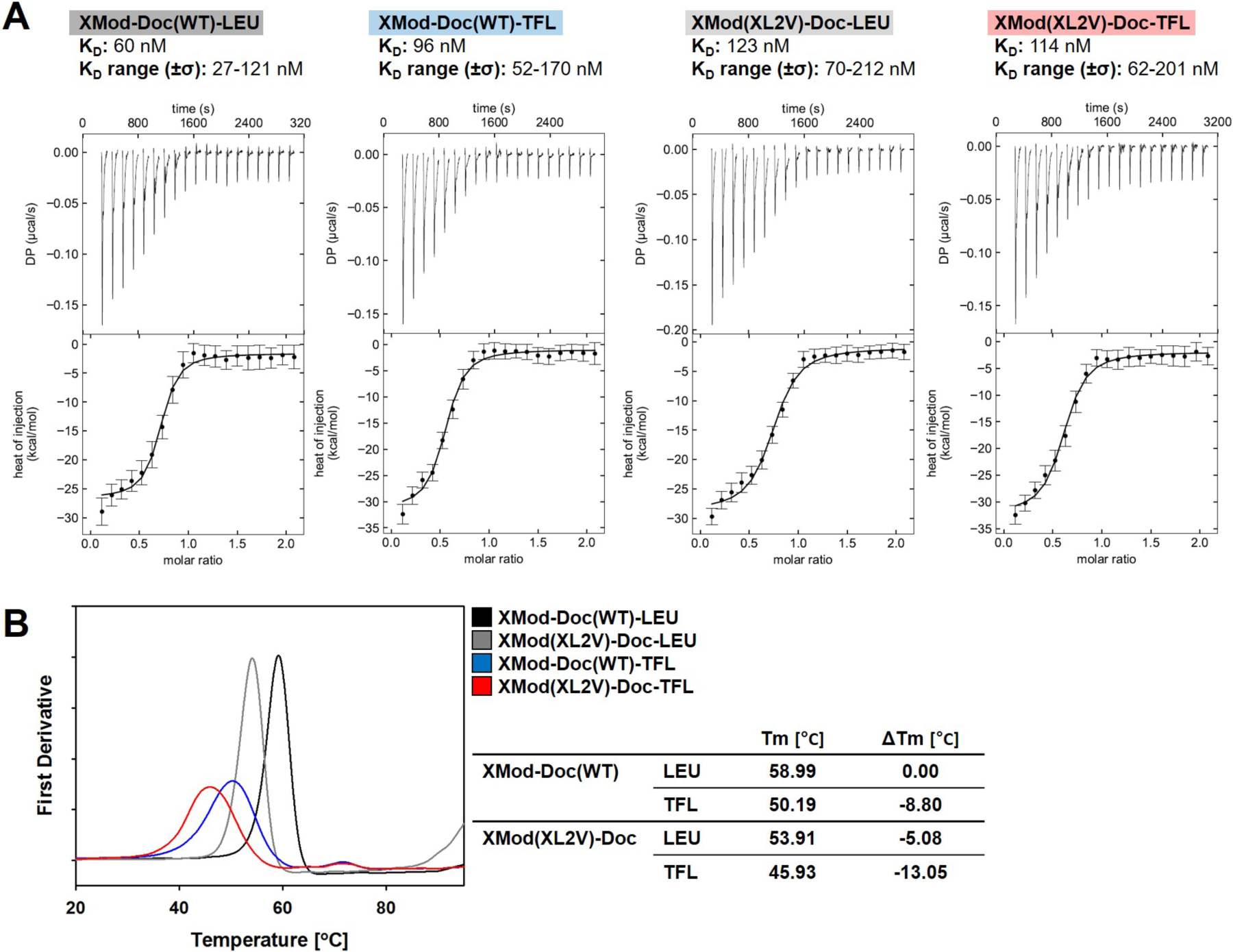
Analysis of bulk biophysical properties of XMod-Doc variants. (A) Binding affinity analyzed by ITC at equilibrium. (B) Thermal melting temperature of XMod-Doc variants measured by DSF.

Finally, we used differential scanning fluorimetry (DSF) to measure thermal denaturation temperatures (**Figure 6B**). DSF analysis showed that mutation of Leu to Val in XMod(XL2V)-Doc under LEU incorporation/expression decreased the thermal melting temperature by ∼5 °C as compared with XMod-Doc(WT)-LEU. Additionally, TFL incorporation decreased thermal denaturation temperatures of both XMod-Doc(WT)-TFL and XMod(XL2V)-Doc-TFL by ∼8 °C. This significant decrease in thermal stability for XMod(XL2V)-Doc-TFL is in contrast to the increase in mechano-stability of XMod that we observed under TFL incorporation (**Figure 5**), and confirms that non-equilibrium mechanical stability need not necessarily be correlated with enhanced thermal stability.

## CONCLUSIONS

Fluorination of proteins is used to tune biophysical properties, and its effects are well studied for systems at equilibrium. Here, for the first time we report the influence of fluorination on a molecular system under non-equilibrium mechanical tension. We investigated the influence of TFL-incorporation on single-molecule unbinding and unfolding reactions within a mechano-stable XMod-Doc:Coh adhesion complex. We designed and produced LEU to VAL mutant variants which minimally disturbed native function of protein, and allowed for TFL to be incorporated globally or only in Doc. Analysis of these variants using equilibrium biophysical stability assays and single-molecule AFM revealed several competing and counterintuitive effects.

Firstly, we found that global fluorination of XMod-Doc(WT) changed the energy landscape of the Doc:Coh unbinding and XMod unfolding reactions. Fluorination tended to rigidify WT complexes, providing a steeper loading rate dependency (lower Δx) for both P1 high-force and P2 low-force rupture events. Secondly, global fluorination of XMod-Doc(WT) generated a new unbinding pathway (P3). We observed a clearly distinguishable rupture class that lacked XMod unfolding (**Figure 3F**) but ruptured at intermediate force. We attributed emergence of this new pathway to alteration of the energy landscape by chirality or hydrophobicity of TFL. To gain more insight, we performed additional measurements at pH 5.5 and found similar ratios between P1, P2 and P3 rupture events (**Figure S6, Table S1**). These dissociation pathways are therefore independent of pH across the range tested (5.5 - 7.2). Thirdly and most significantly, we found that when fluorination was localized to Doc, XMod was mechanically stabilized (**Figure 5**). We consider this a counterintuitive finding given that the same sample was significantly less thermally stable (**Figure 6B**) and had little change in binding affinity (**Figure 6A**).

Taken together, these findings demonstrate how fluorination can modulate folding energy landscapes by lengthening or shortening binding/interaction potentials, generating new unbinding pathways, or even mechanically stabilize adjacent non-fluorinated domains. These features highlight the orthogonality of mechanical and thermodynamic stability and provide new insights into the influence of fluorination on protein stability and function. With this work, we have provided a first look into how fluorination can regulate the mechanical properties of protein complexes and more generally protein-based biomaterials.

## ACKNOWLEDGMENTS

This work was supported by the University of Basel, ETH Zurich, an ERC Starting Grant (MMA-715207), the SNF NCCR in Molecular Systems Engineering, and the Swiss National Science Foundation (Project 200021_175478).

## References

(1) Wang, L.; Xie, J.; Schultz, P. G. Expanding the Genetic Code. Annu. Rev. Biophys. Biomol. Struct. 2006, 35, 225–249.

(2) Johnson, J. A.; Lu, Y. Y.; Van Deventer, J. A.; Tirrell, D. A. Residue-Specific Incorporation of Non-Canonical Amino Acids into Proteins: Recent Developments and Applications. Curr. Opin. Chem. Biol. 2010, 14 (6), 774–780.

(3) Lang, K.; Chin, J. W. Cellular Incorporation of Unnatural Amino Acids and Bioorthogonal Labeling of Proteins. Chem. Rev. 2014, 114 (9), 4764–4806.

(4) Mei, H.; Han, J.; Klika, K. D.; Izawa, K.; Sato, T.; Meanwell, N. A.; Soloshonok, V. A. Applications of Fluorine-Containing Amino Acids for Drug Design. Eur. J. Med. Chem. 2020, 186, 111826.

(5) Vulpetti, A.; Dalvit, C.Fluorine Local Environment: From Screening to Drug Design. Drug Discov. Today 2012, 17 (15-16), 890–897.

(6) Purser, S.; Moore, P. R.; Swallow, S.; Gouverneur, V.Fluorine in Medicinal Chemistry. Chem. Soc. Rev. 2008, 37 (2), 320–330.

(7) Robalo, J. R.; Huhmann, S.; Koksch, B.; Vila Verde, A.The Multiple Origins of the Hydrophobicity of Fluorinated Apolar Amino Acids. Chem 2017, 3 (5), 881–897.

(8) Hoffmann, W.; Langenhan, J.; Huhmann, S.; Moschner, J.; Chang, R.; Accorsi, M.; Seo, J.; Rademann, J.; Meijer, G.; Koksch, B.; Bowers, M. T.; von Helden, G.; Pagel, K.An Intrinsic Hydrophobicity Scale for Amino Acids and Its Application to Fluorinated Compounds. Angew. Chem. Int. Ed Engl. 2019, 58 (24), 8216–8220.

(9) Robalo, J. R.; Vila Verde, A.Unexpected Trends in the Hydrophobicity of Fluorinated Amino Acids Reflect Competing Changes in Polarity and Conformation. Phys. Chem. Chem. Phys. 2019, 21 (4), 2029–2038.

(10) Buer, B. C.; Marsh, E. N. G. Fluorine: A New Element in Protein Design. Protein Sci. 2012, 21 (4), 453–462.

(11) Panchenko, T.; Zhu, W. W.; Montclare, J. K. Influence of Global Fluorination on Chloramphenicol Acetyltransferase Activity and Stability. Biotechnol. Bioeng. 2006, 94 (5), 921–930.

(12) Holzberger, B.; Rubini, M.; Möller, H. M.; Marx, A.A Highly Active DNA Polymerase with a Fluorous Core. Angew. Chem. Int. Ed Engl. 2010, 49 (7), 1324–1327.

(13) Berger, A. A.; Völler, J.-S.; Budisa, N.; Koksch, B.Deciphering the Fluorine Code1 The Many Hats Fluorine Wears in a Protein Environment. Acc. Chem. Res. 2017, 50 (9), 2093–2103.

(14) Salwiczek, M.; Nyakatura, E. K.; Gerling, U. I. M.; Ye, S.; Koksch, B.Fluorinated Amino Acids: Compatibility with Native Protein Structures and Effects on Protein–protein Interactions. Chem. Soc. Rev. 2012, 41 (6), 2135–2171.

(15) Merkel, L.; Budisa, N.Organic Fluorine as a Polypeptide Building Element: In Vivo Expression of Fluorinated Peptides, Proteins and Proteomes. Org. Biomol. Chem. 2012, 10 (36), 7241–7261.

(16) Montclare, J. K.; Tirrell, D. A. Evolving Proteins of Novel Composition. Angew. Chem. Int. Ed Engl. 2006, 45 (27), 4518–4521.

(17) Tang, Y.; Ghirlanda, G.; Petka, W. A.; Nakajima, T.; DeGrado, W. F.; Tirrell, D. A. Fluorinated Coiled-Coil Proteins Prepared in Vivo Display Enhanced Thermal and Chemical Stability. Angew. Chem. Int. Ed. 2001, 40 (8), 1494–1496.

(18) Tang, Y.; Tirrell, D. A. Biosynthesis of a Highly Stable Coiled-Coil Protein Containing Hexafluoroleucine in an Engineered Bacterial Host. J. Am. Chem. Soc. 2001, 123 (44), 11089–11090.

(19) Tang, H.-C.; Lin, Y.-J.; Horng, J.-C. Modulating the Folding Stability and Ligand Binding Affinity of Pin1 WW Domain by Proline Ring Puckering. Proteins 2014, 82 (1), 67–76.

(20) Yoo, T. H.; Link, A. J.; Tirrell, D. A. Evolution of a Fluorinated Green Fluorescent Protein. Proc. Natl. Acad. Sci. U. S. A. 2007, 104 (35), 13887–13890.

(21) Dominguez, M. A., Jr; Thornton, K. C.; Melendez, M. G.; Dupureur, C. M. Differential Effects of Isomeric Incorporation of Fluorophenylalanines into PvuII Endonuclease. Proteins 2001, 45 (1), 55–61.

(22) Parsons, J. F.; Xiao, G.; Gilliland, G. L.; Armstrong, R. N. Enzymes Harboring Unnatural Amino Acids: Mechanistic and Structural Analysis of the Enhanced Catalytic Activity of a Glutathione Transferase Containing 5-Fluorotryptophan. Biochemistry 1998, 37 (18), 6286–6294.

(23) Meng, H.; Kumar, K.Antimicrobial Activity and Protease Stability of Peptides Containing Fluorinated Amino Acids. J. Am. Chem. Soc. 2007, 129 (50), 15615–15622.

(24) Meng, H.; Krishnaji, S. T.; Beinborn, M.; Kumar, K.Influence of Selective Fluorination on the Biological Activity and Proteolytic Stability of Glucagon-like Peptide-1. J. Med. Chem. 2008, 51 (22), 7303–7307.

(25) Horng, J.-C.; Raleigh, D. P. Phi-Values beyond the Ribosomally Encoded Amino Acids: Kinetic and Thermodynamic Consequences of Incorporating Trifluoromethyl Amino Acids in a Globular Protein. J. Am. Chem. Soc. 2003, 125 (31), 9286–9287.

(26) Infante, E.; Stannard, A.; Board, Stephanie J.; Rico-Lastres, P.; Rostkova, E.; Beedle, A. E. M.; Lezamiz, A.; Wang, Y. J.; Gulaidi Breen, S.; Panagaki, F.; Sundar Rajan, V.; Shanahan, C.; Roca-Cusachs, P.; Garcia-Manyes, S.The Mechanical Stability of Proteins Regulates Their Translocation Rate into the Cell Nucleus. Nat. Phys. 2019, 15 (9), 973–981.

(27) Rivas-Pardo, J. A.; Li, Y.; Mártonfalvi, Z.; Tapia-Rojo, R.; Unger, A.; Fernández-Trasancos, Á.; Herrero-Galán, E.; Velázquez-Carreras, D.; Fernández, J. M.; Linke, W. A.; Alegre-Cebollada, J.A HaloTag-TEV Genetic Cassette for Mechanical Phenotyping of Proteins from Tissues. Nat. Commun. 2020, 11 (1), 2060.

(28) Ott, W.; Jobst, M. A.; Schoeler, C.; Gaub, H. E.; Nash, M. A. Single-Molecule Force Spectroscopy on Polyproteins and Receptor–ligand Complexes: The Current Toolbox. J. Struct. Biol. 2016, 197 (1), 3–12.

(29) Chen, Y.; Radford, S. E.; Brockwell, D. J. Force-Induced Remodelling of Proteins and Their Complexes. Curr. Opin. Struct. Biol. 2015, 30, 89–99.

(30) Schönfelder, J.; De Sancho, D.; Perez-Jimenez, R.The Power of Force: Insights into the Protein Folding Process Using Single-Molecule Force Spectroscopy. J. Mol. Biol. 2016, 428 (21), 4245–4257.

(31) Nathwani, B.; Shih, W. M.; Wong, W. P. Force Spectroscopy and Beyond: Innovations and Opportunities. Biophys. J. 2018, 115 (12), 2279–2285.

(32) Jacobson, D. R.; Uyetake, L.; Perkins, T. T. Membrane-Protein Unfolding Intermediates Detected with Enhanced Precision Using a Zigzag Force Ramp. Biophysical Journal 2019.

(33) Yu, H.; Siewny, M. G. W.; Edwards, D. T.; Sanders, A. W.; Perkins, T. T. Hidden Dynamics in the Unfolding of Individual Bacteriorhodopsin Proteins. Science 2017, 355 (6328), 945–950.

(34) Dietz, H.; Berkemeier, F.; Bertz, M.; Rief, M.Anisotropic Deformation Response of Single Protein Molecules. Proc. Natl. Acad. Sci. U. S. A. 2006, 103 (34), 12724–12728.

(35) Sedlak, S. M.; Schendel, L. C.; Gaub, H. E.; Bernardi, R. C. Streptavidin/biotin: Tethering Geometry Defines Unbinding Mechanics. Sci Adv 2020, 6 (13), eaay5999.

(36) Sedlak, S. M.; Schendel, L. C.; Melo, M. C. R.; Pippig, D. A.; Luthey-Schulten, Z.; Gaub, H. E.; Bernardi, R. C. Direction Matters: Monovalent Streptavidin/Biotin Complex under Load. Nano Lett. 2019, 19 (6), 3415–3421.

(37) Carrion-Vazquez, M.; Li, H.; Lu, H.; Marszalek, P. E.; Oberhauser, A. F.; Fernandez, J. M. The Mechanical Stability of Ubiquitin Is Linkage Dependent. Nat. Struct. Biol. 2003, 10 (9), 738–743.

(38) Brockwell, D. J.; Paci, E.; Zinober, R. C.; Beddard, G. S.; Olmsted, P. D.; Smith, D. A.; Perham, R. N.; Radford, S. E. Pulling Geometry Defines the Mechanical Resistance of a Beta-Sheet Protein. Nat. Struct. Biol. 2003, 10 (9), 731–737.

(39) Jobst, M. A.; Milles, L. F.; Schoeler, C.; Ott, W.; Fried, D. B.; Bayer, E. A.; Gaub, H. E.; Nash, M. A. Resolving Dual Binding Conformations of Cellulosome Cohesin-Dockerin Complexes Using Single-Molecule Force Spectroscopy. Elife 2015, 4. https://doi.org/10.7554/eLife.10319.

(40) Nash, M. A.; Smith, S. P.; Fontes, C. M.; Bayer, E. A. Single versus Dual-Binding Conformations in Cellulosomal Cohesin–dockerin Complexes. Curr. Opin. Struct. Biol. 2016, 40, 89–96.

(41) Bogan, A. A.; Thorn, K. S. Anatomy of Hot Spots in Protein Interfaces. J. Mol. Biol. 1998, 280 (1), 1–9.

(42) Voronov-Goldman, M.; Yaniv, O.; Gul, O.; Yoffe, H.; Salama-Alber, O.; Slutzki, M.; Levy-Assaraf, M.; Jindou, S.; Shimon, L. J. W.; Borovok, I.; Bayer, E. A.; Lamed, R.; Frolow, F.Standalone Cohesin as a Molecular Shuttle in Cellulosome Assembly. FEBS Lett. 2015, 589 (14), 1569–1576.

(43) Schoeler, C.; Malinowska, K. H.; Bernardi, R. C.; Milles, L. F.; Jobst, M. A.; Durner, E.; Ott, W.; Fried, D. B.; Bayer, E. A.; Schulten, K.; Gaub, H. E.; Nash, M. A. Ultrastable Cellulosome-Adhesion Complex Tightens under Load. Nat. Commun. 2014, 5, 5635.

(44) Schoeler, C.; Bernardi, R. C.; Malinowska, K. H.; Durner, E.; Ott, W.; Bayer, E. A.; Schulten, K.; Nash, M. A.; Gaub, H. E. Mapping Mechanical Force Propagation through Biomolecular Complexes. Nano Lett. 2015, 15 (11), 7370–7376.

(45) Bernardi, R. C.; Durner, E.; Schoeler, C.; Malinowska, K. H.; Carvalho, B. G.; Bayer, E. A.; Luthey-Schulten, Z.; Gaub, H. E.; Nash, M. A. Mechanisms of Nanonewton Mechanostability in a Protein Complex Revealed by Molecular Dynamics Simulations and Single-Molecule Force Spectroscopy. J. Am. Chem. Soc. 2019, 141 (37), 14752–14763.

(46) Verdorfer, T.; Bernardi, R. C.; Meinhold, A.; Ott, W.; Luthey-Schulten, Z.; Nash, M. A.; Gaub, H. E. Combining in Vitro and in Silico Single-Molecule Force Spectroscopy to Characterize and Tune Cellulosomal Scaffoldin Mechanics. J. Am. Chem. Soc. 2017, 139 (49), 17841–17852.

(47) Liu, Z.; Liu, H.; Vera, A. M.; Bernardi, R. C.; Tinnefeld, P.High Force Catch Bond Mechanism of Bacterial Adhesion in the Human Gut. bioRxiv 2020.

(48) Jindou, S.; Brulc, J. M.; Levy-Assaraf, M.; Rincon, M. T.; Flint, H. J.; Berg, M. E.; Wilson, M. K.; White, B. A.; Bayer, E. A.; Lamed, R.; Borovok, I.Cellulosome Gene Cluster Analysis for Gauging the Diversity of the Ruminal Cellulolytic Bacterium Ruminococcus Flavefaciens. FEMS Microbiol. Lett. 2008, 285 (2), 188–194.

(49) Ding, S. Y.; Rincon, M. T.; Lamed, R.; Martin, J. C. Cellulosomal Scaffoldin-Like Proteins fromRuminococcus Flavefaciens. Journal of 2001.

(50) Salama-Alber, O.; Jobby, M. K.; Chitayat, S.Atypical Cohesin-Dockerin Complex Responsible for Cell Surface Attachment of Cellulosomal Components BINDING FIDELITY, PROMISCUITY, AND STRUCTURAL …. Journal of Biological 2013.

(51) Montclare, J. K.; Son, S.; Clark, G. A.; Kumar, K.; Tirrell, D. A. Biosynthesis and Stability of Coiled-Coil Peptides Containing (2 S, 4 R)-5,5,5-Trifluoroleucine and (2 S, 4 S)-5,5,5-Trifluoroleucine. Chembiochem 2009, 10 (1), 84–86.

(52) Liu, H.; Ta, D. T.; Nash, M. A. Mechanical Polyprotein Assembly Using Sfp and Sortase-Mediated Domain Oligomerization for Single-Molecule Studies. Small Methods 2018, 2 (6), 1800039.

(53) Liu, H.; Schittny, V.; Nash, M. A. Removal of a Conserved Disulfide Bond Does Not Compromise Mechanical Stability of a VHH Antibody Complex. Nano Lett. 2019, 19 (8), 5524–5529.

(54) Yang, B.; Liu, Z.; Liu, H.; Nash, M. A. Next Generation Methods for Single-Molecule Force Spectroscopy on Polyproteins and Receptor-Ligand Complexes. Frontiers in Molecular Biosciences 2020, 7, 85.

(55) Wieduwild, R.; Howarth, M.Assembling and Decorating Hyaluronan Hydrogels with Twin Protein Superglues to Mimic Cell-Cell Interactions. Biomaterials 2018, 180, 253–264.

(56) Zakeri, B.; Howarth, M.Spontaneous Intermolecular Amide Bond Formation between Side Chains for Irreversible Peptide Targeting. J. Am. Chem. Soc. 2010, 132 (13), 4526–4527.

(57) Zakeri, B.; Fierer, J. O.; Celik, E.; Chittock, E. C.; Schwarz-Linek, U.; Moy, V. T.; Howarth, M.Peptide Tag Forming a Rapid Covalent Bond to a Protein, through Engineering a Bacterial Adhesin. Proc. Natl. Acad. Sci. U. S. A. 2012, 109 (12), E690–E697.

(58) Schwaiger, I.; Kardinal, A.; Schleicher, M.; Noegel, A. A.; Rief, M.A Mechanical Unfolding Intermediate in an Actin-Crosslinking Protein. Nat. Struct. Mol. Biol. 2004, 11 (1), 81–85.

(59) Ott, W.; Jobst, M. A.; Bauer, M. S.; Durner, E.; Milles, L. F.; Nash, M. A.; Gaub, H. E. Elastin-like Polypeptide Linkers for Single-Molecule Force Spectroscopy. ACS Nano 2017, 11 (6), 6346–6354.

(60) Yin, J.; Straight, P. D.; McLoughlin, S. M.; Zhou, Z.; Lin, A. J.; Golan, D. E.; Kelleher, N. L.; Kolter, R.; Walsh, C.Genetically Encoded Short Peptide Tag for Versatile Protein Labeling by Sfp Phosphopantetheinyl Transferase. Proc. Natl. Acad. Sci. U. S. A. 2005, 102 (44), 15815–15820.

(61) Baba, T.; Ara, T.; Hasegawa, M.; Takai, Y.; Okumura, Y.; Baba, M.; Datsenko, K. A.; Tomita, M.; Wanner, B. L.; Mori, H.Construction of Escherichia Coli K-12 in-Frame, Single-Gene Knockout Mutants: The Keio Collection. Mol. Syst. Biol. 2006, 2 (1).

(62) Yang, B.; Ayyadurai, N.; Yun, H.; Choi, Y. S.; Hwang, B. H.; Huang, J.; Lu, Q.; Zeng, H.; Cha, H. J. In Vivo Residue-Specific Dopa-Incorporated Engineered Mussel Bioglue with Enhanced Adhesion and Water Resistance. Angew. Chem. Int. Ed. 2014, 53 (49), 13360–13364.

(63) Evans, E.; Ritchie, K.Dynamic Strength of Molecular Adhesion Bonds. Biophys. J. 1997, 72 (4), 1541–1555.

(64) Merkel, R.; Nassoy, P.; Leung, A.; Ritchie, K.; Evans, E.Energy Landscapes of Receptor–ligand Bonds Explored with Dynamic Force Spectroscopy. Nature 1999, 397 (6714), 50–53.

(65) Dudko, O. K.; Hummer, G.; Szabo, A. Theory, Analysis, and Interpretation of Single-Molecule Force Spectroscopy Experiments. Proc. Natl. Acad. Sci. U. S. A. 2008, 105 (41), 15755–15760.

(66) Dudko, O. K. Decoding the Mechanical Fingerprints of Biomolecules. Q. Rev. Biophys. 2016, 49, e3.

(67) Brockwell, D. J.; Beddard, G. S.; Paci, E.; West, D. K.; Olmsted, P. D.; Smith, D. A.; Radford, S. E. Mechanically Unfolding the Small, Topologically Simple Protein L. Biophys. J. 2005, 89 (1), 506–519.

(68) Schoeler, C.; Verdorfer, T.; Gaub, H. E.; Nash, M. A. Biasing Effects of Receptor-Ligand Complexes on Protein-Unfolding Statistics. Phys Rev E 2016, 94 (4-1), 042412.

