## Supplementary Information for "Influence of Fluorination on Single-Molecule Unfolding and Rupture Pathways of a Mechanostable Protein Adhesion Complex"

### Materials & Methods

***Plasmids available on addgene:***

Addgene plasmid #157672: pET28a-Coh-FLN-ELP(MV7E2)3-His-ybbR
Addgene plasmid #157673: pQE80L-Coh-His-SpyTag
Addgene plasmid #157674: pET28a-ybbR-His-ELP(MV7E2)3-FLN-SpyCatcher
Addgene plasmid #157675: pQE80L-His-SpyTag-XMod-Doc(WT)
Addgene plasmid #157676: pQE80L-His-SpyTag-XMod(XL2V)-Doc

***Cloning of Coh-FLN-ELP-His-ybbR***

The gene encoding Coh (type III Cohesin from *Ruminococcus flavefaciens)* was amplified via PCR with two primers: 5'-GA AAT AAT TTT GTT TAA CTT TAA GAA GGA GAT ATA CAT ATG GCG CTC ACA GAC AGA GGA A-3' and 5'-CC GGG TCA GCA GAA CCG GAT CC AGA ACC GGA GCC TGG CTC ACC AGC CT-3'. This PCR-amplified fragment and the backbone from digestion of pET28a-FLN-ELP(MV7E2)3-His-ybbR (pET28a vector harboring FLN, ELP linker, his6x tag, and ybbR tag) with NdeI and BamHI were assembled into a new vector pET28a-Coh-FLN-ELP(MV7E2)3-His-ybbR using Gibson assembly master mix (New England BioLabs) and confirmed by DNA sequence analysis.

***Cloning of Coh-His-SpyTag***

The gene encoding Coh was amplified via PCR with two primers: 5'-CACGTCGACGCCCATATTGTCGCGCTCACAGACAGAGGAAT-3' and 5'-GGCGTCGAGCAGCCCGCCGTGATGGTGATGGTGATGTGG-3'. pQE80 backbone was also amplified via PCR with two primers: 5'-GGCGGGCTGCTCGACG-3' and 5'-GACAATATGGGCGTCGACGTGATG-3' using plasmid pQE80L-SpyTag-ELP-SpyTag, a gift from Mark Howarth (Addgene plasmid # 112634 ; http://n2t.net/addgene:112634 ; RRID:Addgene_112634).[55] Then, those PCR products were assembled into a new vector pQE80L-Coh-His-SpyTag using Gibson assembly master mix and confirmed by DNA sequence analysis.

***Cloning of ybbR-His-ELP-FLN-SpyCatcher***

The SpyCatcher sequence (5'-Overlap region/GS linker-SpyCatcher-Stop codon-Overlap region-3') was chemically synthesized based on the *E. coli* codon usage (GeneArt, Thermo Fisher Scientific). pET28a backbone was also amplified via PCR with two primers: 5'- GATCCGGCTGCTAACAAAGCCCGAAAGGAAGCTGAG-3' and 5'-GGAACCAGAACCGGAGCCCGGAGCCGGTTTAACGGTAAC-3' using plasmid pET28a-ybbR-His-ELP($MV_7E_2$)$_3$-FLN (pET28a vector harboring FLN, ELP linker, his6x tag, and ybbR tag). Then, the synthesized gene string and amplified PCR product were assembled into a new vector pET28a-ybbR-His-ELP(MV7E2)3-FLN-SpyCatcher using Gibson assembly master mix and confirmed by DNA sequence analysis.

***Cloning of His-SpyTag-XMod-Doc(WT), (XL2V), and (DL2V)***

DNA sequences of XMod-Doc(WT) (wild type, type III from *Ruminococcus flavefacien*), and mutant sequences XMod(XL2V)-Doc (X-module: Leu to Val), and XMod-Doc(DL2V) (Dockerin: Leu to Val), were optimized based on *E. coli* codon usage. Then, DNA constructs 5'-Overlap region-Optimized DNA sequences of XMod-Doc variants-Overlap region-3' were chemically synthesized (GeneArt, Thermo Fisher Scientific). The backbone was amplified via PCR using plasmid pQE80L-SpyTag-ELP-SpyTag (Addgene #112634)[55] and two primers: 5'-GTGGCCGTCGAGCTTCG-3' and 5'-TAAGGATCCGGCTGCTAACA-3'. This PCR-amplified backbone and the chemically synthesized fragment were assembled into new vectors pQE80L-His-SpyTag-XMod-Doc(WT), (XL2V), and (DL2V) using Gibson assembly master mix and were confirmed by DNA sequence analysis.

***Expression of proteins***

The constructed recombinant plasmids pET28a-Coh-FLN-ELP(MV7E2)3-His-ybbR and pET28a-ybbR-His-ELP(MV7E2)3-FLN-SpyCatcher were transformed into *E. coli* BL21 (DE3) strain and recombinant plasmid pQE80L-Coh-His-SpyTag, pQE80L-His-SpyTag-XMod-Doc(WT), (XL2V), and (DL2V) were transformed into *E. coli* JW5807 leucine auxotrophic strain (Keio collection). Cells were cultured in 5 ml of Luria-Bertani (LB) medium with 50 µg $ml^{-1}$ kanamycin at 37 °C overnight. The culture was transferred to 100 mL of Terrific broth (TB) medium with 50 µg $ml^{-1}$ kanamycin and cultivated at 37 °C and 200 rpm until an optical density at 600 nm ($OD_{600}$) of ~0.8-1.0 was reached. Recombinant protein expression was induced upon addition of 0.5 mM isopropyl-β-D-thio-galactopyranoside (IPTG) and the culture was further incubated at 20 °C and 200 rpm for 12 h. The cells were harvested by centrifugation at 4,000 g for 20 min at 4 °C. The cell pellets were stored at -80 °C until further purification.

***Residue-specific TFL incorporation***

The *Escherichia coli* JW5807 leucine auxotroph (Keio collection) was used as a host strain for the expression of TFL-incorporated XMod-Doc variants. The recombinant vector pQE80L-His-SpyTag-XMod-Doc(WT), (XL2V), and (DL2V) were transformed into JW5807 leucine auxotroph, and a single colony was picked and cultured in 3 ml of Luria-Bertani (LB) medium with 100 µg $mL^{-1}$ ampicillin at 37 °C overnight. The culture was transferred to 10 mL of M9 minimal medium (100 mL of M9 salt (67.8 g of $Na_2HPO_4$, 30.0 g of $KH_2PO_4$, 5.0 g of NaCl, and 10.0 g of $NH_4Cl$ in 1 L distilled water (DW)), 20 mL of 20% glucose, 1 mL of 2 M $MgSO_4$, 1 mL of 0.1 M $CaCl_2$, and 1 mg of thiamine-hydrochloric acid in 1 L DW) with 20 canonical amino acids (40 mg $L^{-1}$), including leucine, 100 µg $mL^{-1}$ ampicillin, and 50 µg $mL^{-1}$ kanamycin, and cultured at 37 °C overnight. The culture was transferred to 50 mL of M9 minimal medium with 19 canonical amino acids, a limited amount of leucine (0.09 mM), 100 µg $mL^{-1}$ ampicillin and 50 µg $mL^{-1}$ kanamycin was cultivated at 37 °C and 180 rpm for approximately 7 h to reach the stationary phase with an optical density at 600 nm ($OD_{600}$) of ~0.9-1.0. After confirmation of the stationary phase, TFL (5,5,5-Trifluoro-DL-leucine, Sigma-Aldrich) was introduced at a concentration of 2 mM in the medium, and the expression of TFL-incorporated XMod-Doc(WT), (XL2V), and (DL2V) were induced by the addition of 1 mM IPTG. The culture was further incubated at 37 °C and 180 rpm for 6 h. The cells were harvested by centrifugation at 4,000 g for 20 min at 4 °C. The cell pellets were stored at -80 °C until further purification.

***Protein purification***

All expressed recombinant proteins including TFL-incorporated XMod-Doc variants include a hexa-histidine ($His_6$) tag to enable purification using immobilized metal affinity chromatography. The harvested cell pellet was resuspended in a lysis buffer (50 mM Tris, 50 mM NaCl, 0.1% Triton X-100, 5 mM $MgCl_2$; pH 8.0), and disrupted with a sonic dismembrator. The lysate was centrifuged at 14,000 g for 20 min at 4 °C. The supernatant was collected and incubated with Ni-NTA resin (Thermo Fisher Scientific), loaded onto a column, washed with wash buffer (TBS buffer (25 mM Tris and 72 mM NaCl) with 25 mM imidazole; pH 7.2), and eluted in elution buffer (TBS buffer with 500 mM imidazole; pH 7.2). The eluted protein solution was buffer-exchanged to TBS buffer pH 7.2 using AKTA and HiPrep column, further purified by Superose SEC column, and finally stored in 33% glycerol at -20 °C.

***Size-exclusion chromatography***

Coh and XMod-Doc(WT), (XL2V), and (DL2V) mutants binding assay was performed using size exclusion chromatography in TBS $Ca^{2+}$ buffer (25 mM Tris, 72 mM NaCl, 1 mM $CaCl_2$; pH 7.2). Each sample was injected into a pre-equilibrated Superose 12 10/300 GL column. Binding behavior was confirmed by comparing elution time between mixture of Coh(Coh-FLN-ELP-His-ybbR) and XMod-Doc(WT)-LEU or (XL2V)-LEU vs Coh only, and between mixture of Coh(Coh-His-SpyTag) and XMod-Doc(DL2V)-LEU vs XMod-Doc(DL2V)-LEU only (Figure S4).

***High-resolution mass spectrometry***

Purified XMod-Doc(WT)-LEU and -TFL, and XMod-Doc(XL2V)-LEU and -TFL protein solution were desalted using Zeba™ spin desalting column (Thermo Fisher Scientific) and diluted to a concentration of 0.2-1.0 mg $mL^{-1}$ with a final concentration of 0.1% formic acid. The separation of the sample protein analytes was carried out using an UltiMate™ 3000 UHPLC-system equipped with 50 mm Phenomenex Jupiter C4 column (Thermo Fischer Scientific) with a diameter of 2.0 mm, 300Å pore size, and 5 µm particle size. 1µL of protein solution was injected for all analyses and the column was kept at 30 °C. HRMS-spectra were acquired on a Bruker maXis 4G ESI-QTOF (Bruker Daltonics) and data deconvolution was done with Bruker Compass DataAnalysis 4.4. TFL incorporation yield was calculated based on the integrated mass spectrum of each peak.

***Surface preparation and protein immobilization for AFM-SMFS***

The surface modification of cantilever and coverglasses and the protein immobilization were done in the same manner as previously illustrated (Figure 1H). Cantilevers were cleaned by UV-ozone treatment for 40 min and coverglasses were soaked in piranha etching solution and rinsed with distilled water (DW). Then, cantilevers and coverglasses were treated with 3-Aminopropyl (diethoxy) methylsilane (APDMES, ABCR GmbH, Karlsruhe, Germany) to silanize the surfaces with amine groups. The amine groups subsequently reacted to a NHS group from sulfosuccinimidyl 4-(N-maleimidomethyl)cyclohexane-1-carboxylate (sulfo-SMCC; Thermo Fischer Scientific) in 50 mM HEPES buffer pH 7.5 for 30 min. The thiol group from Coenzyme A (CoA, 200 µM) reacted to a maleimide group from sulfo-SMCC in coupling buffer (50 mM sodium phosphate, 50 mM NaCl, 10 mM EDTA, pH 7.2) for 2 hrs. Finally, the ybbR-tagged proteins Coh-FLN-ELP-His-ybbR and pre-conjugated ybbR-His-ELP-FLN-SpyCatcher-SpyTag-XMod-Doc variants were site-specifically anchored to the surface using SFP-mediated ligation to CoA in $Mg^{2+}$ supplemented TBS buffer (25 mM Tris, 72 mM NaCl, 10 mM $MgCl_2$, pH 7.2). This resulted in covalent immobilization of Coh and XMod-Doc variants to cantilever and cover glasses, respectively. Protein-immobilized cantilevers and coverglasses were extensively washed and kept in TBS $Ca^{2+}$ buffer (25 mM Tris, 72 mM NaCl, 1 mM $CaCl_2$; pH 7.2) prior to immediate use. SpyTag-SpyCatcher conjugation of ybbR-His-ELP-FLN-SpyCatcher and His-SpyTag-XMod-Doc variants were done by mixing two proteins with same molar ratio in TBS buffer and pre-incubation for 1 hr prior to ybbR tag ligation.

***AFM-SMFS measurement and data analysis***

Force spectroscopy measurements with Coh and XMod-Doc variants were conducted in the same manner as previously illustrated using automated AFM-based SMFS (Force Robot 300, JPK Instruments). SMFS data were recorded in TBS $Ca^{2+}$ buffer (25 mM Tris, 72 mM NaCl, 1 mM $CaCl_2$; pH 7.2) at room temperature with constant pulling speeds of 400 nm $s^{-1}$, 800 nm $s^{-1}$, 1600 nm $s^{-1}$ and 3200 nm $s^{-1}$. Total 139,218 force-extension curves were acquired. Force-extension curves were filtered and analyzed by a combination of software available on the AFM instrument and custom python scripts. The majority of data traces contained no interactions, non-specific interaction, or complex multiplicity of interactions. Therefore, the data traces were filtered by searching for contour length increments that matched the lengths of the fingerprint domains, FLN (≈36 nm). Theoretical contour length increment was calculated based on the equation $\Delta L_c$ = (0.365 nm/AA) × (# AAs in POI) - $L_f$, where $\Delta L_c$ is expected contour length increment and $L_f$ is end-to-end length of folded protein domain. For FLN, $\Delta L_c$ = 36.9 nm - $L_f$, where $L_f$ is typically < 5nm. The total number of force-extension curves matching this criterion was 371 out of 22,747, 670 out of 22,153, 1,117 out of 45,787, and 1,021 out of 48,531 for XMod-Doc(WT)-LEU, -TFL, XMod-Doc(XL2V)-LEU, and –TFL, respectively.

For dynamic force spectra, the rupture or unfolding forces vs. loading rate was plotted and median forces and loading rates for each pulling speed were fitted to Bell-Evans model[63,64] to estimate the effective distance to the transition state (Δx) and the intrinsic dissociation rate or unfolding rate ($k_{off}$) in the absence of force. Data were fitted using the Dudko-Hummer-Szabo model[65,66] with a cusp-like barrier ($\nu = 0.5$) to estimate Δx, $k_{off}$, and energy barrier height (ΔG).

For direct comparison with the same cantilever, both XMod(XL2V)-Doc-LEU and -TFL were immobilized on different areas of the same surface and a single cantilever was used to probe each spot using constant speed pulling at 100 nm $s^{-1}$. After 500 approach-retraction cycles at one area, the surface was moved to the other area. This cycle was been done twice for each XMod(XL2V)-Doc-LEU and -TFL. P-values were calculated using data sets between LEU and TFL incorporated XMod-Doc variants with two-tailed t-test and indicated as n.s. $p \geq 0.01$, *$p < 0.01$, **$p < 0.001$, and ***$p < 0.0001$.

***ITC***

The binding affinity between Coh and XMod-Doc variants was measured using the MicroCal ITC-200 instrument (Malvern Instruments). Protein concentrations were chosen to give data suitable for accurate $K_D$ determination. The cell volume was 200 µL and each injection volume was 2 µL, with 20 injections. Each 5 µM solution of XMod-Doc variants ((WT)-LEU, -TFL, and (XL2V)-LEU, -TFL) was transferred into the sample cell. 50 µM solution of Coh (Coh-His-SpyTag) in the injection syringe was added to the cell and the data was recorded as the power needed to maintain the reference cell and sample cell at the same temperature. All the measurements were conducted in TBS buffer (25 mM Tris, 72 mM NaCl, 1 mM $CaCl_2$; pH 7.2) and solutions were degassed under a vacuum prior to the experiments. The data from titrations were analyzed and the calculated dissociation constants are summarized in Figure 5A.

***Nano DSF***

The XMod-Doc variants were loaded into capillaries and subjected to be analyzed by nano differential scanning fluorimetry (nanoDSF) using a Prometheus NT 48 (NanoTemper Technologies). The protein thermal unfolding curves were determined in a range between 20 °C and 95 °C with a linear thermal ramp of 1 °C/min. The melting temperatures (Tm) were determined from the change in a tryptophan fluorescence ratio 350/330 nm and calculated based on the maximum point of the first derivative of thermal denaturation curve. Protein samples were prepared at 30 µM in TBS $Ca^{2+}$ buffer (25 mM Tris, 72 mM NaCl, 1 mM $CaCl_2$; pH 7.2). All samples were measured in triplicate.

***Protein sequences***

**His-SpyTag-XMod-Doc(WT)**

MKGSSHHHHHHVDAHIVMVDAYKPTKLDGHNTVTSAVKTQYVEIESVDGFYFNTEDKFDTAQIKKAVLHTVYNEGYTGDDGVAVVLREYESEPVDITAELTFGDATPANTYKAVENKFDYEIPVYYNNATLKDAEGNDATVTVYIGLKGDTDLNNIVDGRDATATLTYYAATS

TDGKDATTVALSPSTLVGGNPESVYDDFSAFLSDVKVDAGKELTRFAKKAERLIDGRDASSILTFYTKSSVDQYKDMAANEPNKLWDI
VTGDAEEE

**His-SpyTag-XMod-Doc(XL2V)**

MKGSSHHHHHHVDAHIVMVDAYKPTKLDGHNTVTSAVKTQYVEIESVDGFYFNTEDKFDTAQIKKAVVHTVYNEGYTGDDGVAV
VVREYESEPVDITAEVTFGDATPANTYKAVENKFDYEIPVYYNNATVKDAEGNDATVTVYIGVKGDTDLNNIVDGRDATATLTYYAA
TSTDGKDATTVALSPSTLVGGNPESVYDDFSAFLSDVKVDAGKELTRFAKKAERLIDGRDASSILTFYTKSSVDQYKDMAANEPNKL
WDIVTGDAEEE

**His-SpyTag-XMod-Doc(DL2V)**

MKGSSHHHHHHVDAHIVMVDAYKPTKLDGHNTVTSAVKTQYVEIESVDGFYFNTEDKFDTAQIKKAVLHTVYNEGYTGDDGVAV
VLREYESEPVDITAELTFGDATPANTYKAVENKFDYEIPVYYNNATLKDAEGNDATVTVYIGLKGDTDVNNIVDGRDATATVTYYAAT
STDGKDATTVAVSPSTVVGGNPESVYDDFSAFVSDVKVDAGKEVTRFAKKAERVIDGRDASSIVTFYTKSSVDQYKDMAANEPNKV
WDIVTGDAEEE

**ybbR-His-ELP-FLN-SpyCatcher**

MGTDSLEFIASKLAHHHHHHWGSGHGVGVPGMGVPGVGVPGVGVPGVGVPGVGVPGVGVPGVGVPGVGVPGEGVPGEGVP
GVGVPGMGVPGVGVPGVGVPGVGVPGVGVPGVGVPGVGVPGVGVPGEGVPGEGVPGVGVPGMGVPGVGVPGVGVPGVGV
PGVGVPGVGVPGVGVPGVGVPGEGVPGEGVPGWPSGSADPEKSYAEGPGLDGGECFQPSKFKIHAVDPDGVHRTDGGDGFVVT
IEGPAPVDPVMVDNGDGTYDVEFEPKEAGDYVINLTLDGDNVNGFPKTVTVKPAPGSGSGSGSVDTLSGLSSEQGQSGDMTIEED
SATHIKFSKRDEDGKELAGATMELRDSSGKTISTWISDGQVKDFYLYPGKYTFVETAAPDGYEVATAITFTVNEQGQVTVNGKATKG
DAHI

**Coh-FLN-ELP-His-ybbR**

MALTDRGMTYDLDPKDGSSAATKPVLEVTKKVFDTAADAAGQTVTVEFKVSGAEGKYATTGYHIYWDERLEVVATKTGAYAKKGA
ALEDSSLAKAENNGNGVFVASGADDDFGADGVMWTVELKVPADAKAGDVYPIDVAYQWDPSKGDLFTDNKDSAQGKLMQAYF
FTQGIKSSSNPSTDEYLVKANATYADGYIAIKAGEPGSGSGSGSADPEKSYAEGPGLDGGECFQPSKFKIHAVDPDGVHRTDGGDGF
VVTIEGPAPVDPVMVDNGDGTYDVEFEPKEAGDYVINLTLDGDNVNGFPKTVTVKPAPGSGSGSHGVGVPGMGVPGVGVPGVG
VPGVGVPGVGVPGVGVPGVGVPGVGVPGEGVPGEGVPGVGVPGMGVPGVGVPGVGVPGVGVPGVGVPGVGVPGVGVPGVG
VPGEGVPGEGVPGVGVPGMGVPGVGVPGVGVPGVGVPGVGVPGVGVPGVGVPGVGVPGEGVPGEGVPGWRGHHHHHHGS
DSLEFIASKLA

### Supporting Figures & Tables

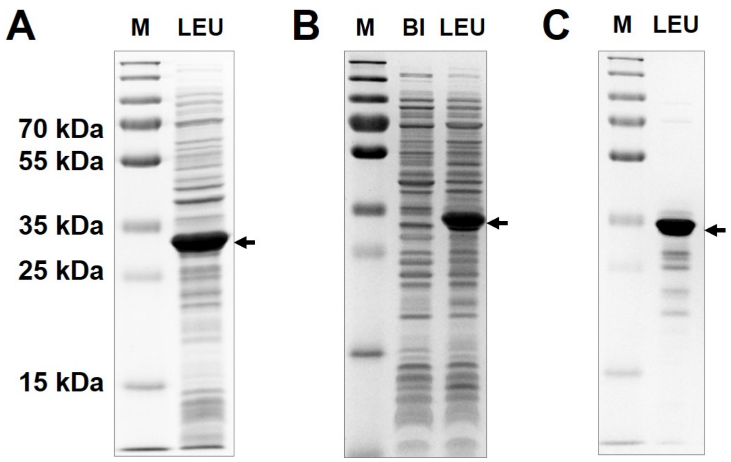


**Figure S1.** Expression of XMod-Doc variants with natural amino acid LEU. SDS-PAGE analysis of successful expression of (A) XMod-Doc(WT)-LEU and (B) XMod(XL2V)-Doc-LEU. (C) SDS-PAGE analysis of successful expression and purification of XMod-Doc(DL2V)-LEU. M: molecular marker, BI: before induction, and LEU: after induction with LEU.

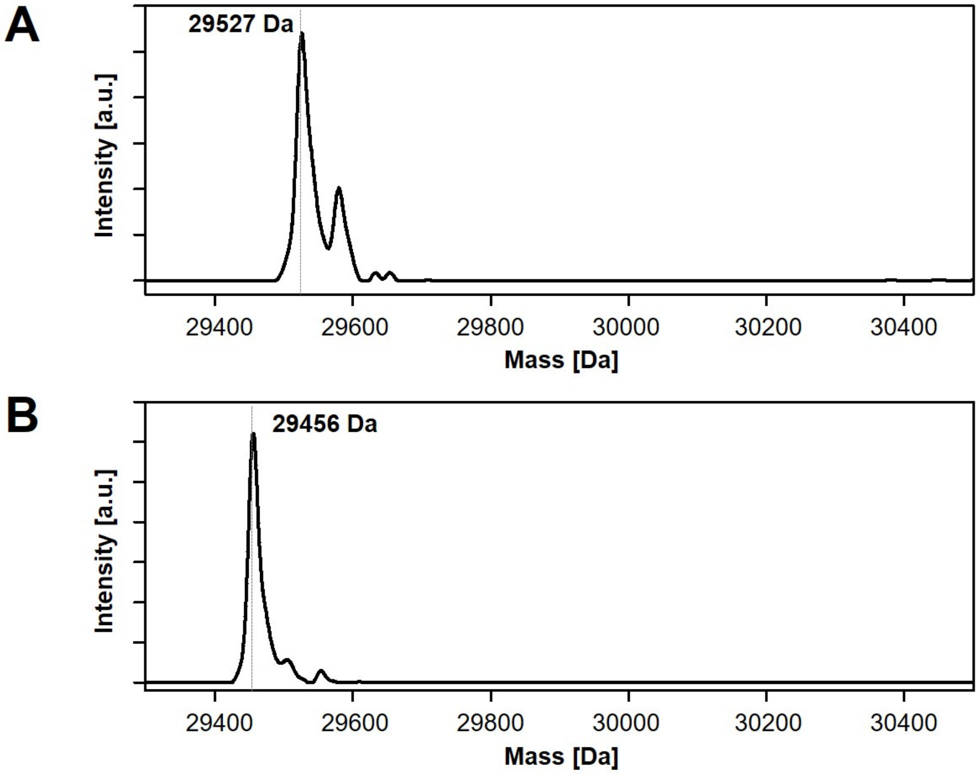


**Figure S2.** HRMS analysis without TFL incorporation. (A) Major mass peak of XMod-Doc(WT)-LEU and (B) XMod(XL2V)-Doc-LEU.

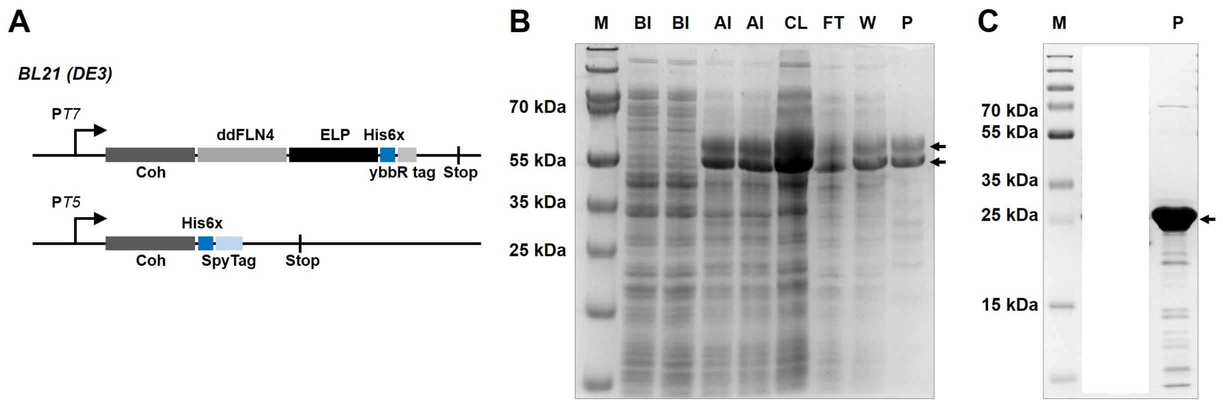


**Figure S3.** Expression of Coh variants. (A) Schematic constructs of plasmid for Coh-FLN-ELP-His-ybbR for AFM-SMFS analysis and Coh-His-SpyTag for ITC analysis. SDS-PAGE analysis of successful expression and purification of (B) Coh-FLN-ELP-His-ybbR and (C) Coh-His-SpyTag. M: molecular marker, BI: before induction, AI: after induction, CL: cleared lysate, FT: flow through, W: wash, and P: purified.

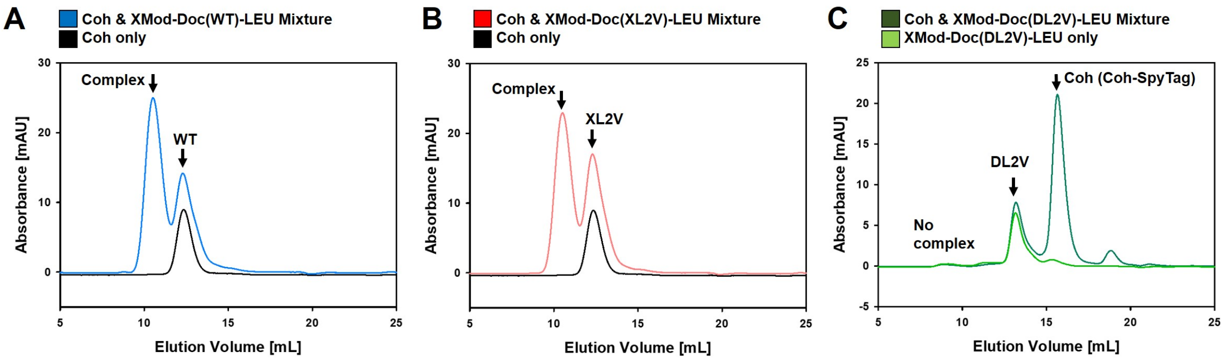


**Figure S4.** Analysis of binding between XMod-Doc varians and Coh with size exclusion chromatography (SEC). Complex formation was confirmed for (A) XMod-Doc(WT)-LEU and (B) XMod-Doc(XL2V)-LEU with Coh-FLN-ELP-His-ybbR protein. (C) Complex formation was not observed between XMod-Doc(DL2V)-LEU and Coh-His-SpyTag protein.

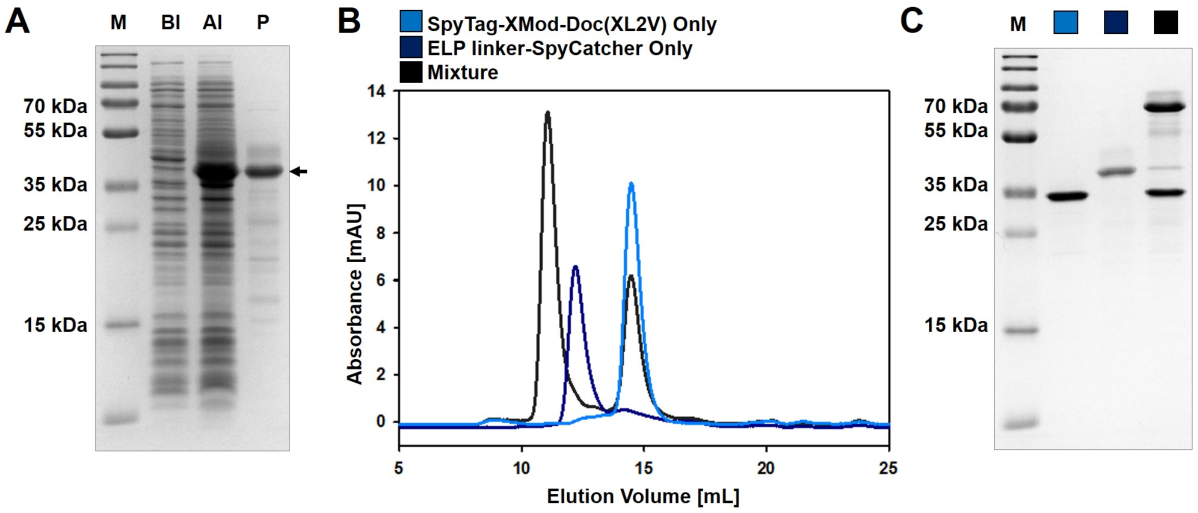


**Figure S5.** Conjugation between SpyTag and SpyCatcher domain. (A) SDS-PAGE analysis of successful expression and purification of ELP linker-SpyCatcher protein. (B) SEC analysis of successful binding/conjugation between SpyTag-XMod-Doc(XL2V)-LEU and ELP linker-SpyCatcher protein. (C) Covalent conjugation by isopeptide bond confirmed by SDS-PAGE analysis. M: molecular marker, BI: before induction, AI: after induction, and P: purified.

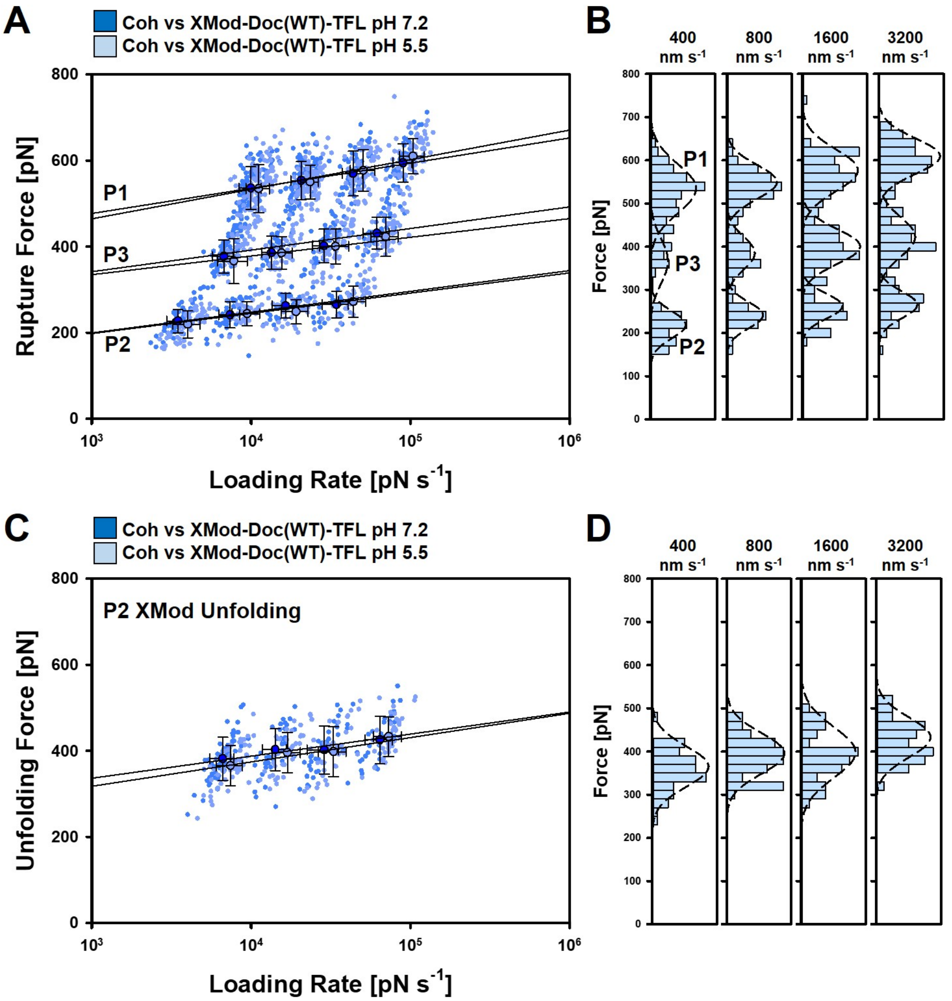


**E**

| | | Coh:XMod-Doc Rupture | | | | | | P2 (XMod unfolding event) | |
|---|---|---|---|---|---|---|---|---|---|
| | | P1 (XMod folded) | | P2 (XMod unfolded) | | P3 (XMod folded) | | | |
| | | Δx (nm) | $k_{off}$ ($s^{-1}$) | Δx (nm) | $k_{off}$ ($s^{-1}$) | Δx (nm) | $k_{off}$ ($s^{-1}$) | Δx (nm) | $k_{unfolding}$ ($s^{-1}$) |
| **XMod-Doc(WT) (pH 5.5)** | TFL | 0.14 | $5.9 \times 10^{-6}$ | 0.20 | $3.1 \times 10^{-3}$ | 0.22 | $1.1 \times 10^{-6}$ | 0.22 | $1.1 \times 10^{-6}$ |

**Figure S6.** AFM-SMFS of XMod-Doc(WT)-TFL at pH 5.5. (A) Dynamic force spectra of XMod-Doc(WT)-TFL:Coh complex rupture forces (skyblue) according to the loading rate using acetate buffer pH 5.5. (B) Histograms of complex rupture forces with different pulling speeds. (C) Dynamic force spectra of XMod unfolding forces according to the loading rate. (D) Histograms of XMod unfolding forces with different pulling speed. Skyblue circles represent the median rupture force/loading rate at each pulling speed of 400, 800, 1600, and 3200 nm $s^{-1}$. Error bars are ±1 s.d. Solid lines are least square fits to the Bell-Evans model. Dynamic force spectra of XMod-Doc(WT)-TFL (blue) at pH 7.2 was adapted from Figure 3. (E) Kinetic parameters from fitting to Bell-Evans model.

**Table S1.** Percentages of rupture classes.

| | | | **Class I (Intact XMod)** | **Class II (XMod unfolded)** | **Class III (Low force with Intact XMod)** |
|---|---|---|---|---|---|
| **XMod-Doc(WT)** | **LEU** | 400 nm/s | 88% (89) | 12% (12) | 0% (0) |
| | | 800 nm/s | 78% (66) | 22% (19) | 0% (0) |
| | | 1600 nm/s | 79% (69) | 21% (18) | 0% (0) |
| | | 3200 nm/s | 72% (71) | 28% (27) | 0% (0) |
| | **TFL** | 400 nm/s | 45% (101) | 26% (58) | 29% (64) |
| | | 800 nm/s | 55% (92) | 23% (38) | 22% (36) |
| | | 1600 nm/s | 41% (60) | 27% (40) | 32% (47) |
| | | 3200 nm/s | 49% (65) | 22% (30) | 29% (39) |
| **XMod-Doc(XL2V)** | **LEU** | 400 nm/s | 71% (193) | 29% (79) | 0% (0) |
| | | 800 nm/s | 67% (171) | 33% (84) | 0% (0) |
| | | 1600 nm/s | 73% (218) | 27% (82) | 0% (0) |
| | | 3200 nm/s | 65% (189) | 35% (101) | 0% (0) |
| | **TFL** | 400 nm/s | 56% (144) | 44% (111) | 0% (0) |
| | | 800 nm/s | 64% (181) | 36% (101) | 0% (0) |
| | | 1600 nm/s | 63% (164) | 37% (95) | 0% (0) |
| | | 3200 nm/s | 57% (129) | 43% (96) | 0% (0) |

| | | | Class I (Intact XMod) | Class II (XMod unfolded) | Class III (Low force with Intact XMod) |
|---|---|---|---|---|---|
| **XMod-Doc(WT)** | **TFL (pH 5.5)** | 400 nm/s | 59% (110) | 27% (51) | 14% (27) |
| | | 800 nm/s | 51% (95) | 22% (41) | 27% (51) |
| | | 1600 nm/s | 42% (73) | 21% (37) | 36% (63) |
| | | 3200 nm/s | 41% (68) | 24% (39) | 35% (58) |
